# PhenoCellPy: A Python package for biological cell behavior modeling

**DOI:** 10.1101/2023.04.12.535625

**Authors:** Juliano F. Gianlupi, T.J. Sego, James P. Sluka, James A. Glazier

## Abstract

PhysiCell, a mechanistic center-based agent-based modeling framework, describes sequences of cell behaviors in a very elegant way with its phenotype submodel. However, its phenotypes are not usable in other frameworks, and the creation of a new phenotype is not straight forward. PhenoCellPy is an open-source Python package that implements PhysiCell’s phenotypes and their methods in a general way, that is easily extended and embedded into other python-based models. PhenoCellPy defines methods for modeling sequences of cell behaviors and states (*e.g.*, the cell cycle, or the Phases of cellular necrosis). PhenoCellPy defines Python classes for the Cell Volume (which it subdivides between the cytoplasm and nucleus) and its evolution, the state of the cell and the behaviors the cell displays in each state (called the Phase), and the sequence of behaviors (called the Phenotype). PhenoCellPy’s can extend existing modeling frameworks as an embedded model. It integrates with the frameworks by defining the cell states (phases), signaling when a state change occurs, if division occurs, and by receiving information from the framework (*e.g.*, how much interferon the cell is exposed to). PhenoCellPy can function with any python-based modeling framework that supports Python 3, NumPy and SciPy.

## 1 Introduction

Repeatedly in biology we see stereotyped sequences of transitions between relatively distinct phases. While such sequences of phases occur at the scale of societies, ecosystems, animals, organelles, macro-molecular machines, the classical example occurs at the scale of cells. This paper will primarily discuss sequences of cell phases, but the concepts and tools developed generalize easily to other biological agents.

A typical example for cells would be the cell cycle, either simplified as proliferation followed by quiescence (see Section 3.2), or in its more complete form of G0/G1 followed by S, G2, M (Figure 1a, see Sections 3.4 and 3.5). Another example would be a model of a SARS-CoV-2 infected cell, the cell transitions from an uninfected state (or phase) to an infected state with no viral release (eclipse phase), then to a virus releasing state, and finally dies [1] (Figure 1b).

**Figure 1:**
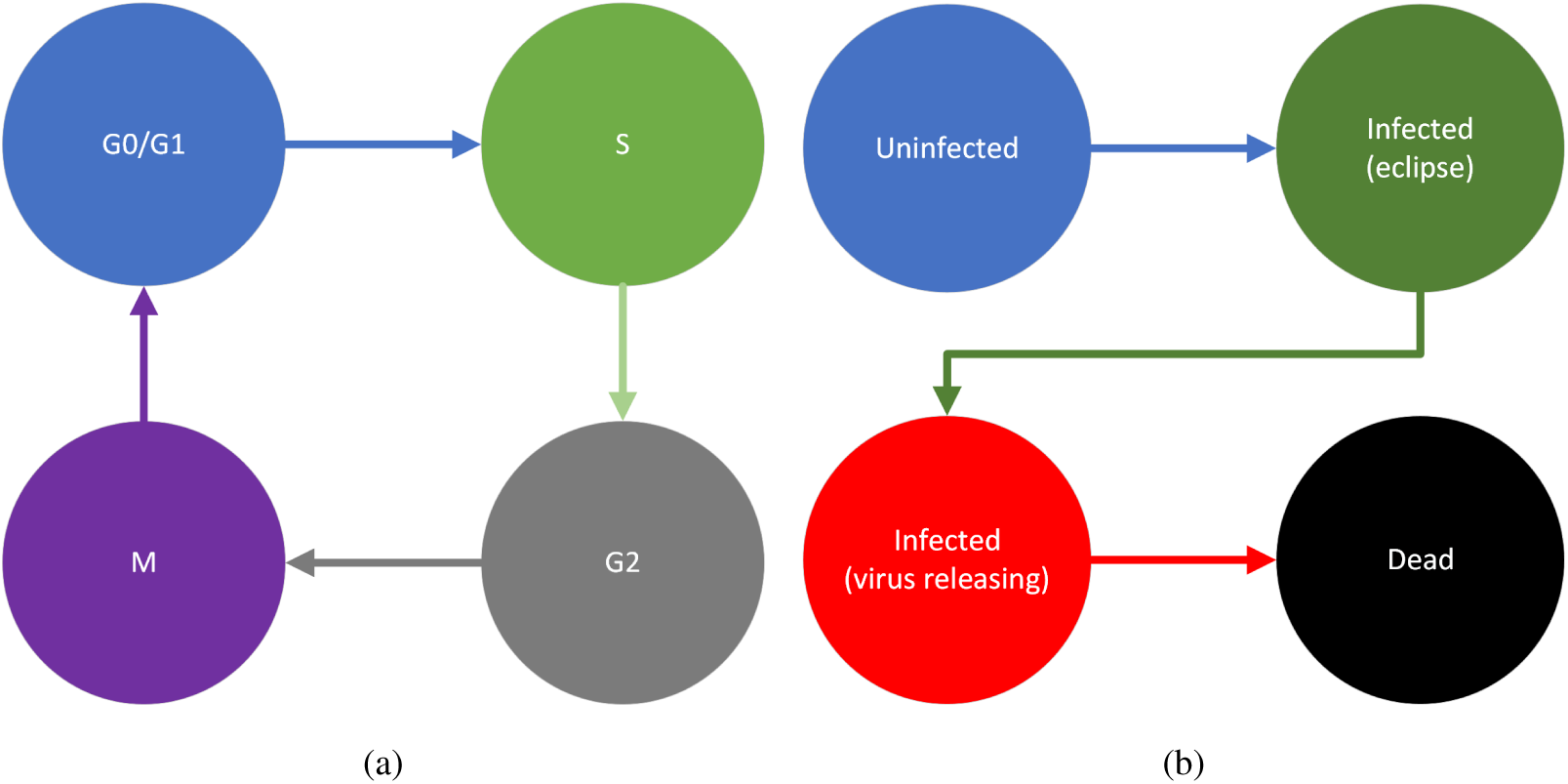
Example sequences of cell behaviors. a) Cell cycle. b) Stages of viral infection of a cell.

For biological systems modeling, it is important to have a structured definition of biological states, sequences of states, and conditions to go from one state to the next. Abstracting cellular behaviors, or, more generally, biological-agent behavior, into a computer model can require the modeler to simultaneously understand the biology of interest and the implementation of that biology in a computational framework. To make modeling more accessible to biologists we need to simplify the transition from the biological description to computational implementation.

There are many computational frameworks for modeling multiscale cellular system: CompuCell3D, Tissue Forge, Morpheus, PhysiCell, Artistoo, FLAME, *etc.* Currently, each modeling framework requires a different type of description of behaviors and behavior transition triggers, and often do not have a defined standard for basic biological *processes* like “cell cycle”. Often implementations of cell, or biological-agent, behaviors are platform-specific and even modeler-specific. This results in models of cell behaviors that are difficult to interpret, share and re-use as well as being restricted to the platform they were created in. This situation makes it difficult to separate the biology being modeled from the framework it is being modelled in. Ideally, the description of the biology being modeled should be separate from its algorithmic implementation. For instance, a model of sequence of cell behaviors built for CompuCell3D [2] of, *e.g.*, infected cell states [1], can’t be simply copied and re-used as-is in some other platform (*e.g.*, Tissue Forge [3]). The modeler has to back out the underlying conceptual model of the phases and their transition rules by interpreting the original implementation, which mixes the biological concepts and the implementing code, and re-code it according to the model specification structure in the new platform. This interpretation is time consuming and subject to many types of error, including misunderstanding of the original model structure and transition rules, mismapping of model-specific parameter values and the possibility of introducing coding errors when the conceptual rules are remapped to code in the new model specification language.

The PhysiCell [4] package has developed a really elegant way to do this. Phenotypes^1^ are defined as the sequence of behaviors and are implemented as a class. The different behaviors that make up a phenotype are called phase. The trigger to go from one phase to the next can be set to either be deterministic after a set time, or stochastic with a set rate. The transition can also depend on environmental factors, or on the cell size. We believe PhysiCell’s approach to phenotypes would be valuable in many other multicellular model frameworks, *e.g.*, in CompuCell3D [2], and Tissue Forge. Currently PhysiCell’s implementation of phenotypes is in C++ and is closely linked with the PhysiCell package, making it difficult to reuse in other modeling frameworks.

PhenoCellPy implements and makes available for general use PhysiCell’s phenotype functionality in in the form of a Python package. It also makes the process of generating new phenotypes, phases, and phase change triggers easy. This addresses the issues of lack of standards and platform specificity, by creating an easy to use and platform-independent Python package that can be embedded in other models. PhenoCellPy is an open-source package, its source code is available at its GitHub repository [5]. Although the concepts and methods of PhenoCellPy are general to many types of biological agents (cells, mitochondria, nucleus, certain organelles) it was built with the cell as the focus.

Phenotype here can mean the cell cycle, the sequential stages of necrosis, the fact that the cell is alive, the fact that it is dead, the different stages of viral infection, *etc*. Phenotype is, then, the set of observable characteristics or traits of an organism.

In PhenoCellPy we create methods and Python classes to define cell behaviors, sequences of cell behaviors, and rules for the behavior switching. We have built several pre-packaged models representing pheno-types of cell behaviors to show-case PhenoCellPy’s capabilities (see Section 3).

An abstract Phenotype consists of one or more Phases (Section 2.2), each Phase defines the volume of the cell, and the volume change rates the Phase displays. It also defines which is the next Phase of the Phenotype, what conditions trigger Phase change, if the agent should divide when exiting the Phase, what behaviors occur immediately on Phase entry and just before Phase exit (*e.g.*, changing the target volume). The cell volume dynamics are handled by the Cell Volume class. The cell volume is subdivided among the solid and fluid cytoplasm, solid and fluid nucleus, and a calcified fraction.

**Listing 1:**
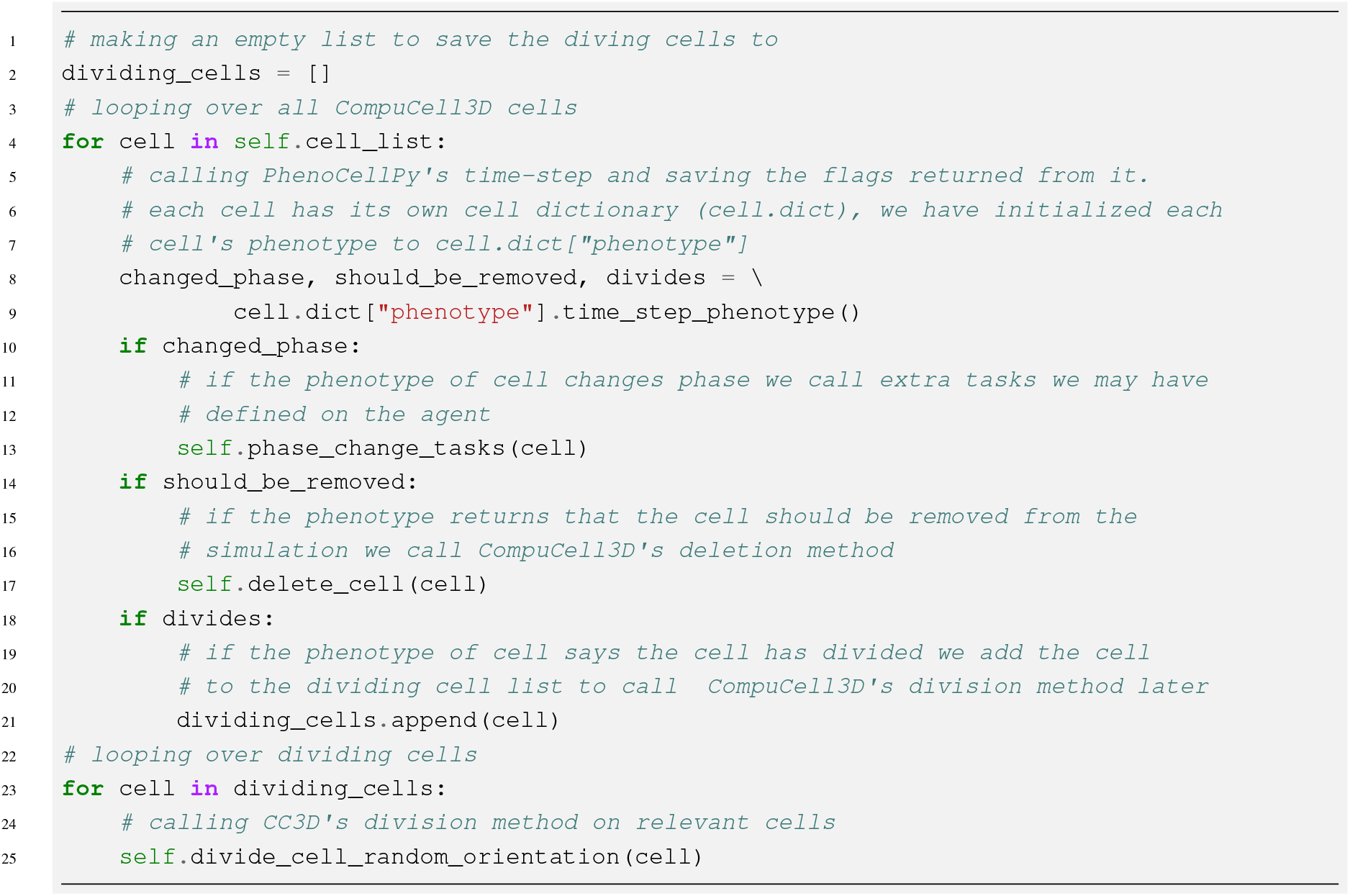
Example implementation of continuous PhenoCellPy tasks in CompuCell3D. Each CC3D cell has its own cell dictionary (`cell.dict`) that can have custom data. We have saved each cell’s Phenotype to the key “phenotype” in the dictionary, see Listing 4.

PhenoCellPy is intended to be used with other python-based modeling frameworks as an embedded model. A PhenoCellPy phenotype should be attached to each relevant agent in the main model, then for each main model agent that has a Phenotype the Phenotype time-step method should be called at every main model time-step. The Phenotype time-step will return boolean flags for cell division, removal from the simulation (*e.g.*, cell death, migration), and cell division, the user is then responsible for performing tasks based on those flags. For instance, in the Necrosis Standard CompuCell3D example (see online), the CompuCell3D cell changes its cell type (see [2] for a definition of cell type in CompuCell3D) when changing from the “hydropic/osmotic swell”^2^ Phase to the “lysed”^3^ Phase (see Section 3.7 for information about the pre-built necrosis Phenotype). Listing 1 shows a generic implementation of continuous PhenoCellPy tasks in a CompuCell3D simulation.

We currently have developed and tested PhenoCellPy embedded models in CompuCell3D [2] (CC3D) and Tissue Forge [3] (TF).

## 2 PhenoCellPy Overview

PhenoCellPy makes the construction of new behavior models easy by breaking them into component parts. The Phenotype class (Section 2.3) is the main “container” of behavior. It can contain all the stages of infection, all the stages of the cell cycle, all the phases of necrosis, *etc*. The Phenotype is broken down into component phases, represented by the Phase class (Section 2.2). The Phase class contains more specific behaviors, *e.g.*, osmotic swelling (necrosis), cell rupture, volume decrease during apoptosis. The Phase class also defines what should be the volume change rates, and what is the trigger for Phase change. Finally the cell volume dynamics are handled by the Cell Volume class (Section 2.1). Figure 2 shows schematics of how PhenoCellPy is organized.

**Figure 2:**
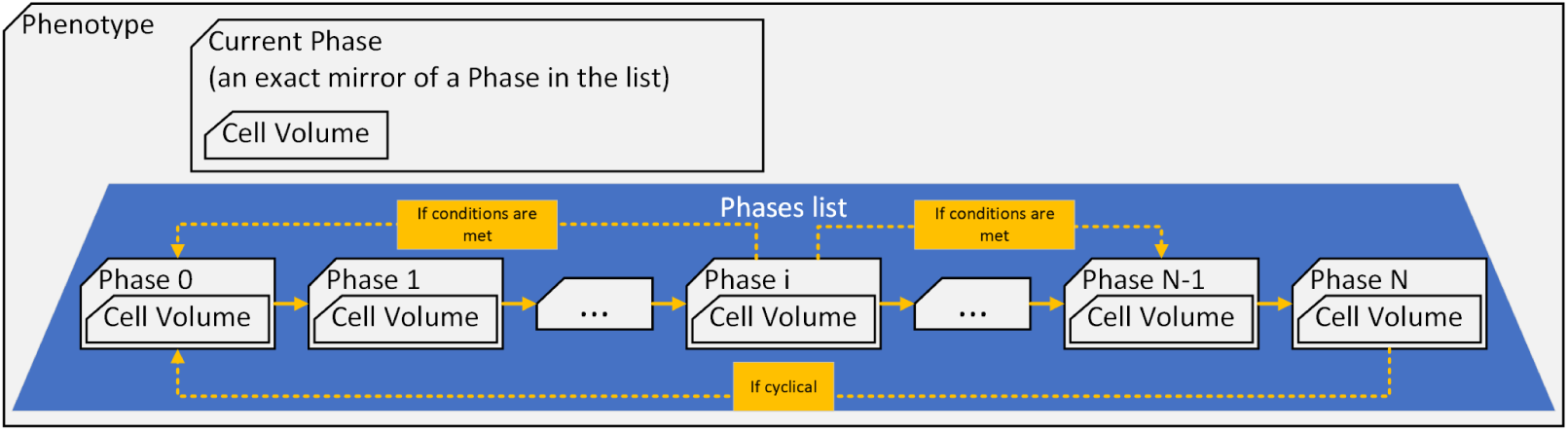
How PhenoCellPy is organized.Gray boxes are PhenoCellPy classes, the blue trapezoid is the lists of constituent Phases. Ownership of objects is conveyed through overlaying shapes (*e.g.*, the Phenotype owns the list of Phases, the Phase owns the Cell Volume). Yellow arrows indicate sequence in the list of Phases. Any Phase could go to any other Phase, as dictated by the modeler. A Phase can have exits to any number of Phases, and a Phase can have entrances from multiple Phases.

We will now briefly describe how to use each class and give an overview of how they work. Starting with the Cell Volume class and working our way up to the Phenotype class. In Section 2.4, we give a brief example of PhenoCellPy’s use. For further detail on how PhenoCellPy is implemented in Python see Supplemental Materials A.

### 2.1 Cell Volume Class

The Cell Volume class defines how big the simulated cell is and how much of its volume is taken by the nucleus and cytoplasm. It also separates the cellular volume into fluid and solid fractions, and has a concept of a calcified volume fraction. All the volumes and fractional volumes we define, as well as volume change rates are in Table 1. The user of PhenoCellPy should decide if the different volumes should be explicitly included in their model, or if they will use only the cytoplasm and nuclear volumes (without a distinction of solid and fluid fraction), or, simply, the total cell volume. The default volumes used by PhenoCellPy are from MCF-7 cells [6] in cubic microns.

**Table 1:** Volumes and volume change rates defined by the Cell Volume class. Volumes marked with ^∗^ are the dynamic volumes.

| Volume type | Symbol | Change Rate | Symbol |
| --- | --- | --- | --- |
| Total | $V$ | Fluid | $r_F$ |
| Total Nuclear | $V_N$ | Nuclear Solid | $r_{NS}$ |
| Nuclear Solid | $V_{NS}^*$ | Cytoplasm Solid | $r_{CS}$ |
| Nuclear Fluid | $V_{NF}$ | Calcified Fraction | $r_C$ |
| Total Cytoplasm | $V_C$ | | |
| Cytoplasm Solid | $V_{CS}^*$ | | |
| Cytoplasm Fluid | $V_{CF}$ | | |
| Cytoplasm to<br>nuclear ratio | $f_{CN}$ | | |
| Total Fluid | $V_F^*$ | | |
| Fluid Fraction | $f_F$ | | |
| Calcified<br>Fraction | $f_C^*$ | | |

The volume dynamics model works by relaxing the dynamic volumes (marked with *^∗^* in Table 1) to their target volumes using their respective volume change rates with a system of ODEs (Equations 1). Then the other volumes are set as relations of the dynamic volumes. It is important to note that, while the volumes and target volumes are attributes of the Cell Volume class, the volume change rates are not, they are an attribute of the Phase class. We made this separation because how fast a cell changes its volume is a property of its state (Phase in PhenoCellPy), therefore the change rates are an attribute of the Phase class. Equations 1 show ODE system governing the dynamic volumes, in those equations the superscript *tg* denotes target. Equations 2 show how the other volumes are calculated from the dynamic volumes and each other.

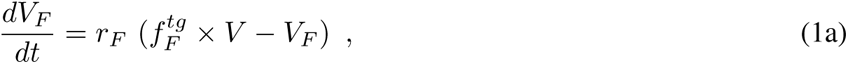

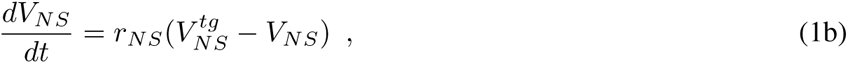

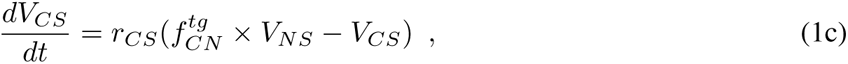

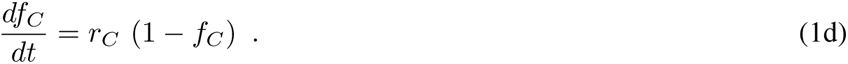

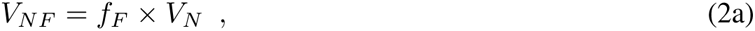

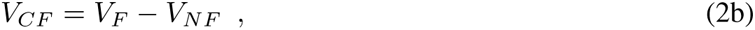

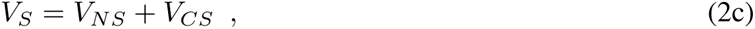

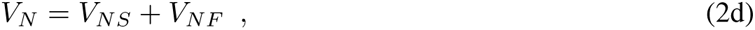

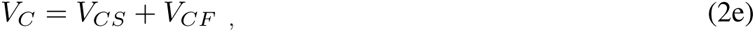

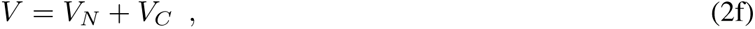

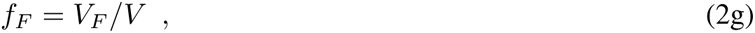

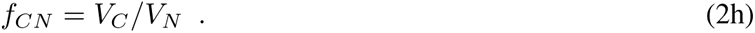

Note that Equation 1d implies that the target calcified fraction is always 1, we are adopting this default behavior from PhysiCell [4]. For most Phases predefined in PhenoCellPy the calcification rate is zero.

Figure 3 shows the Cell Volume class attributes and functions. It also schematizes how the volume update function works. It uses the volumes attributes from the Cell Volume class together with the volume change rates passed to it by the Phase class to update the volumes of the Cell Volume class.

**Figure 3:**
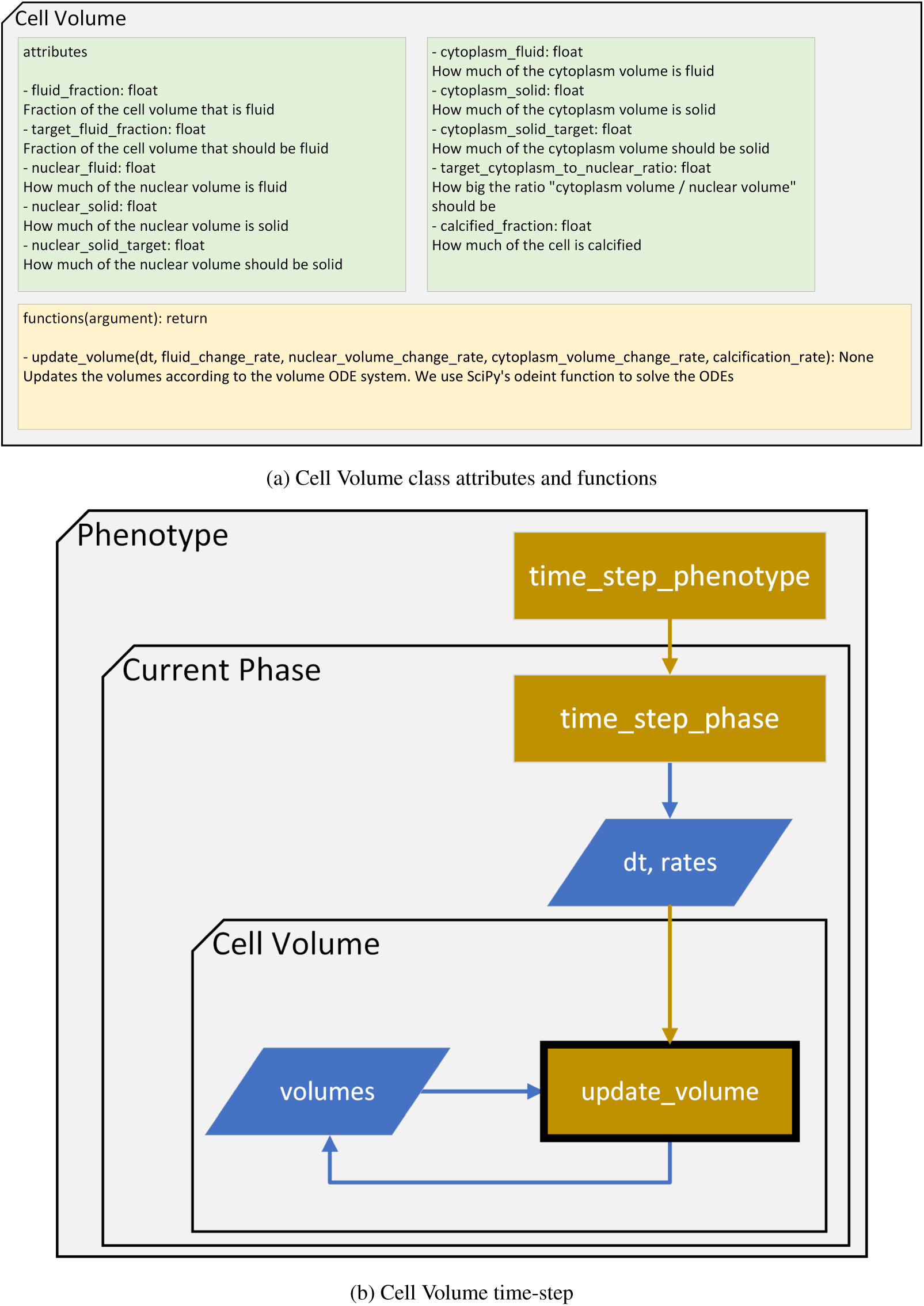
Cell Volume class attributes and functions (3a), and Cell Volume update volume (3b). The Cell Volume update is highlighted by the black outline. Gray boxes are PhenoCellPy classes, yellow rectangles are functions being called, blue parallelograms are information being passed to/from functions, green diamonds are decisions. Yellow arrows mean function call, blue arrows are information being passed.

### 2.2 Phase Class

The phase is the “base unit” of the phenotype. It defines volume change rates, checks for Phase change, and performs phase-specific tasks on phase entry, phase exit and during each time-step.

The phase change check and phase specific tasks are user-definable functions. For flexibility, we impose that all user-definable functions **must** be able to take any number of arguments as inputs (*i.e.*, be a Python *args function). Otherwise, the way PhenoCellPy calls those functions would have to change as the number of arguments changes. The user can define their function is such a way that the *args are unused or unnecessary. Our default transition functions and entry/exit functions functions do not use the args. For PhenoCellPy’s *alpha* version the modeler is responsible for defining, transferring and updating the arguments, creating interface APIs to facilitate this will be included with PhenoCellPy by release.

Examples of phase specific tasks are doubling the target volumes when entering the proliferating Phase (see Sections 3.2 and 3.3), or halving the target volumes after division. In-Phase tasks could be checking if the quantity of a nutrient a cell has access to. Figure 4a shows the Phase class attributes and functions, and Figure 4b the flowchart for the Phase class time-step.

**Figure 4:**
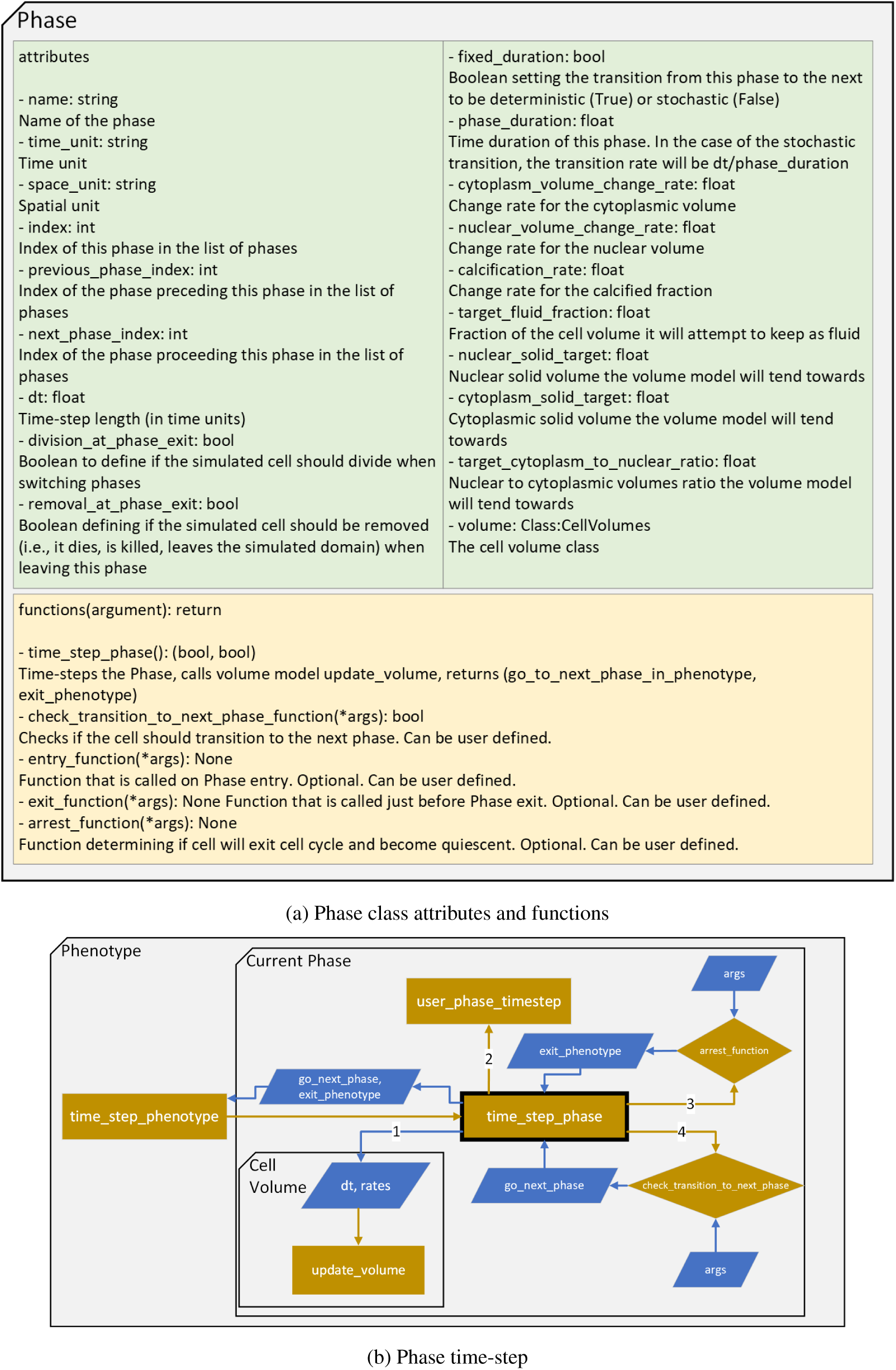
Phase class attributes and functions (4a), and Phase time-step flowchart (4b). The time-step function is highlighted by the black outline, the order in which it performs operations is overlayed on the arrows. Gray boxes are PhenoCellPy classes, yellow rectangles are functions being called, blue parallelograms are information being passed to/from functions, yellow diamonds are decision-making functions. Yellow arrows mean function call, blue arrows are information being passed.

#### 2.2.1 Phase transition

PhenoCellPy has two pre-defined transition functions that are used by our pre-built models, the deterministic phase transition and the stochastic phase transition. For the deterministic transition the Phase evaluates if the time spent in this Phase (*T*) is greater than the Phase period (*τ*), Equation 3. For the stochastic case we draw a probability from a Poisson distribution for a single event occurring (Equation 4). The Poisson probability depends on the time-step length (*dt*) and expected Phase period (*τ*). We cannot approximate 1 *− e^−dt/τ^ ≈ dt/τ* in Equation 4, as we don’t know what values of *dt* and *τ* the modeler will use.

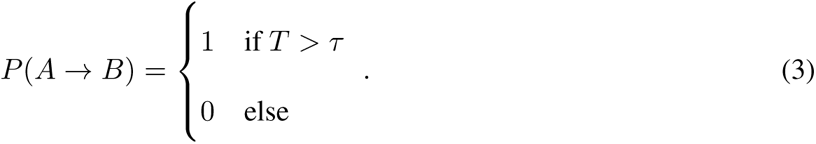

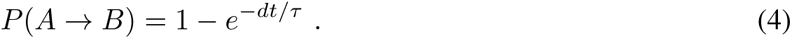

The user can define their own transition functions that may depend on any number of simulation parameters, *e.g.*, oxygen levels, having a neighboring cell, signaling molecules. One of our CompuCell3D [2] examples uses a custom transition function, Ki-67 Basic Cycle Improved Division (Section 4.1.1), to ensure cells don’t divide early. The custom transition function checks that the in-simulation volume of the simulated cell has reached the doubling volume, Equation 6.

If the Phase can transition to several Phases (instead of just one), the user can use the transition function to select which Phase to go to.

### 2.3 Phenotype Class

The Phenotype is the main concept of PhenoCellPy, in PhenoCellPy “Phenotype” means any sequence of distinct cell behaviors. For instance, a quiescent-proliferating cell cycle is a phenotype with two phases (quiescence, and growth/division); the necrotic phenotype starts with a osmotic swelling phase, followed by dissolution of the cell after it bursts.

PhenoCellPy supports cyclical and acyclical Phenotypes, as well as Phenotypes with an arbitrary sequence of Phases. To facilitate a Phenotype end-point we have defined a method to exit the Phenotype cycle and go into a special senescent Phase. To make use of this functionality, the modeler has to define the optional arrest function which is a member of the Phase class (Section 2.2). The arrest function is called by the Phase time-step function (Appendix A.1.2) and its return value is used by the Phenotype time-step function to exit the Phenotype cycle.

The Phenotype class owns the list of all Phases that make it. It switches Phases or goes to the senescent Phase when it receives the respective signals from the Phase time-step, and performs specific user-defined Phenotype tasks each time-step. Figure 5a shows the Phenotype class attributes and functions, and Figure 5b the flowchart for its time-step.

**Figure 5:**
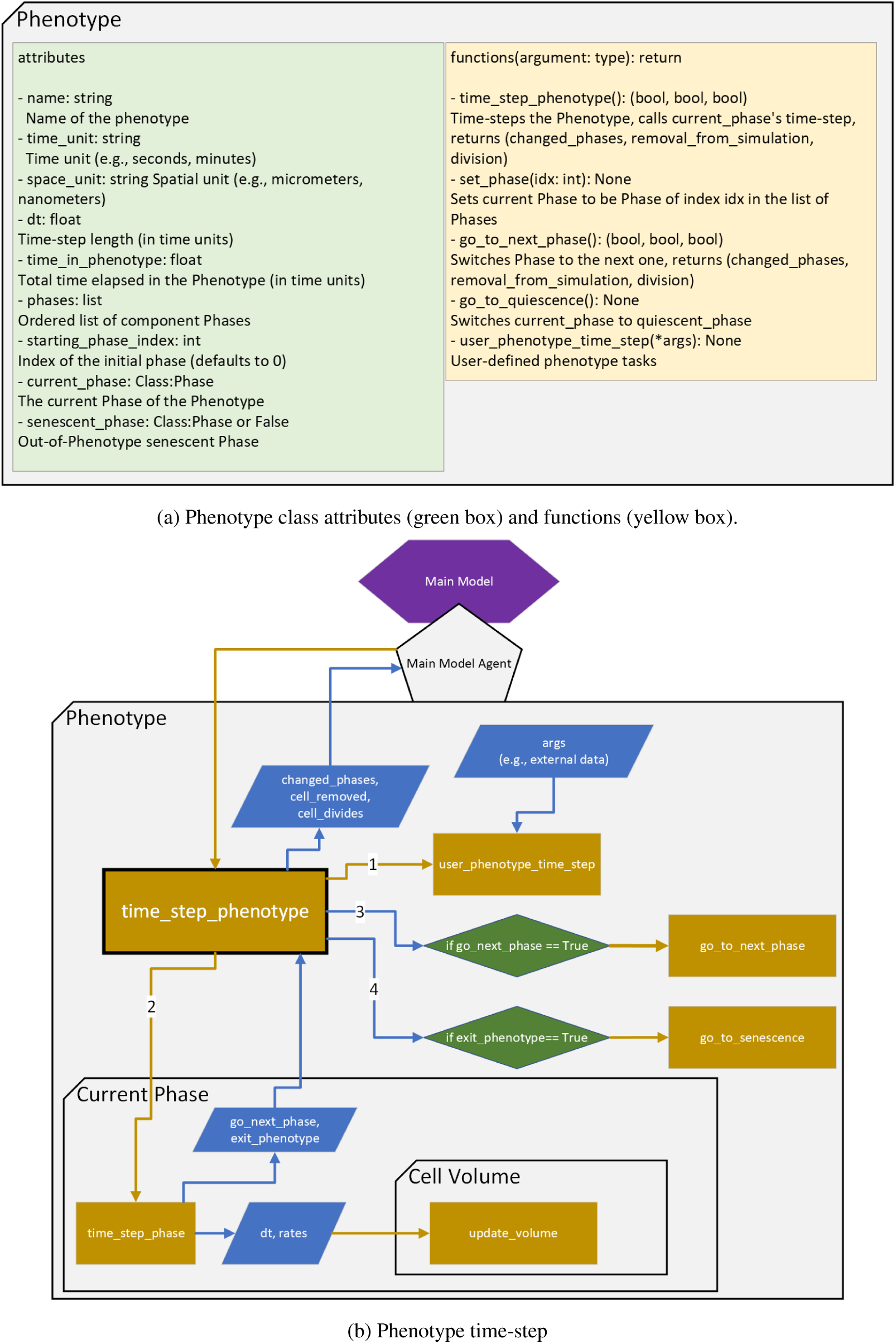
Phenotype class attributes and functions (5a), and Phenotype time-step flowchart (5b). The time-step function is highlighted by the black outline, the order in which it performs operations is overlayed on the arrows. Purple hexagon represents the model PhenoCellPy is embedded in (*i.e.*, the main model), the gray pentagon is an agent from the main model. Gray boxes are PhenoCellPy classes, yellow rectangles are functions being called, blue parallelograms are information being passed to/from functions, green diamonds are decisions. Yellow arrows mean function call, blue arrows are information being passed.

### 2.4 Using PhenoCellPy

PhenoCellPy’s intended use is as an embedded model, meaning it should be loaded into some other modeling platform, *e.g.*, CompuCell3D [2], Tissue Forge [3]. CompuCell3D already supports other modeling frameworks as embedded models, namely SBML [7], Antimony [8], and MaBoSS [9].

To use PhenoCellPy, the modeler has to import PhenoCellPy and should initialize a Phenotype object at the simulation start. The Phenotype initialization (in the version of the package at time of writing) takes as arguments:

- The phenotype name
- Time-step length (*dt*)
- Time and space units
- The list of Phases that make the Phenotype
- Lists for the Phases’ target volumes, initial volumes, volume change rates
- The initial volume of the simulated cell
- The starting Phase index
- The senescent Phase
- User-defined Phenotype time-step tasks and initial arguments for them
- List of user-defined phase time-step tasks and initial arguments for them

All arguments, except the time-step period, for Phenotype initialization are optional. Listing 2 show the import and initialization of a pre-built Phenotype.

**Listing 2:**
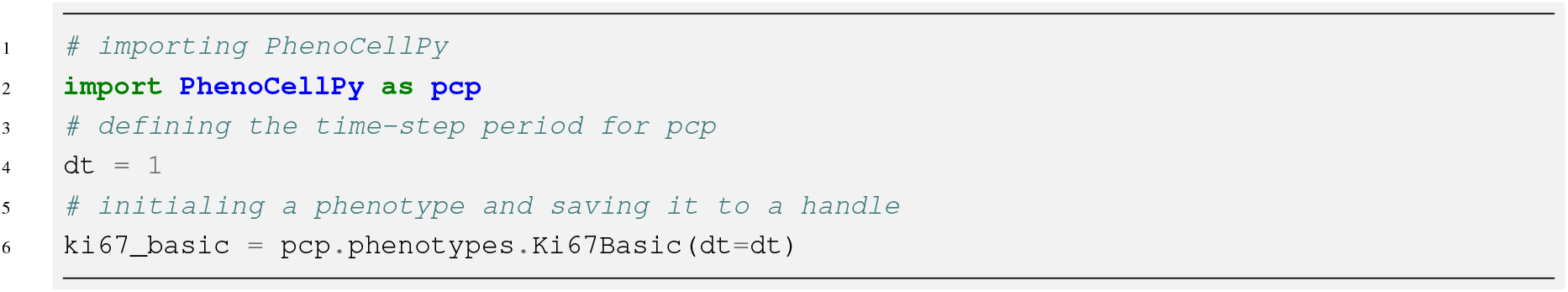
Initialization of a pre-built Phenotype using the default argument values explicitly.

#### 2.4.1 Initialization

A user can also define their own Phenotype, to do so they have to initialize each constituent Phase of the Phenotype and pass them as a list to the Phenotype object initialization. Listing 3, shows an example of a custom Phenotype being built, some of the Phases use custom transition functions. The example in Listing 3 only passes the necessary attributes to the classes.

After having initialized the Phenotype the user has to attach it to each agent that will use that Phenotype. How to do this attachment is modeling platform dependent, in CompuCell3D the recommended method is to make the Phenotype a cell dictionary (Listing 4), we have created an utility function that does this (Listing 5). In Tissue Forge recommended method is to have a dictionary with cell ids as keys and Phenotype as items (Listing 6).

**Listing 3:**
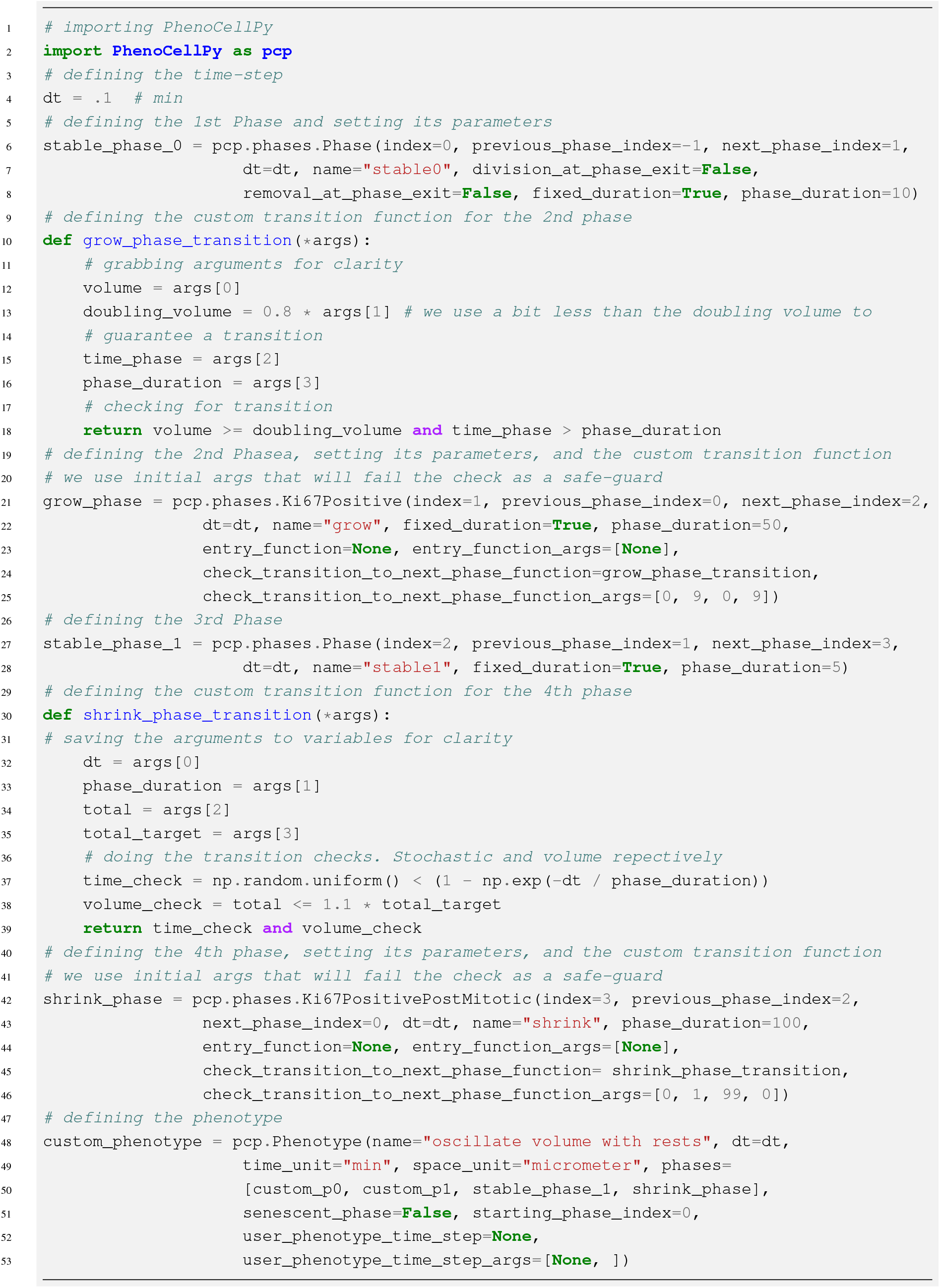
Initialization of a custom Phenotype.

**Listing 4:**
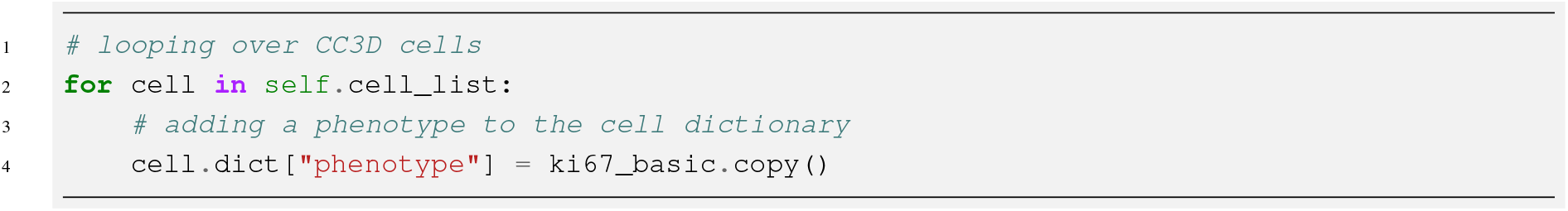
Attaching a Phenotype to a CompuCell3D cell. Listing 2 shows the phenotype initialization.

**Listing 5:**
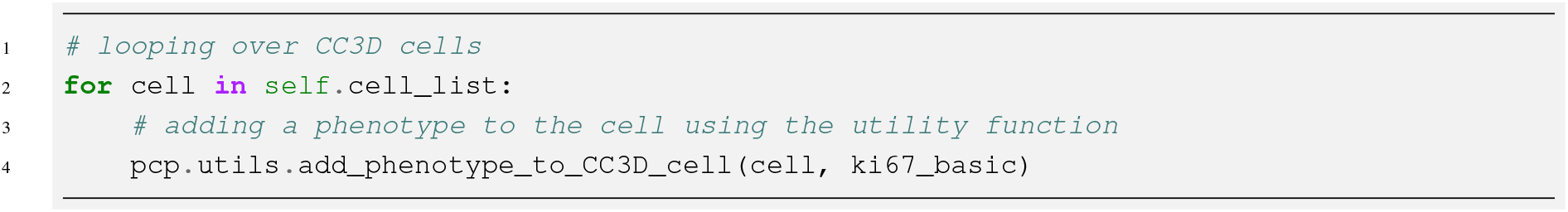
Attaching a Phenotype to a CompuCell3D cell using PhenoCellPy’s utility function. Listing 2 shows the phenotype initialization.

**Listing 6:**
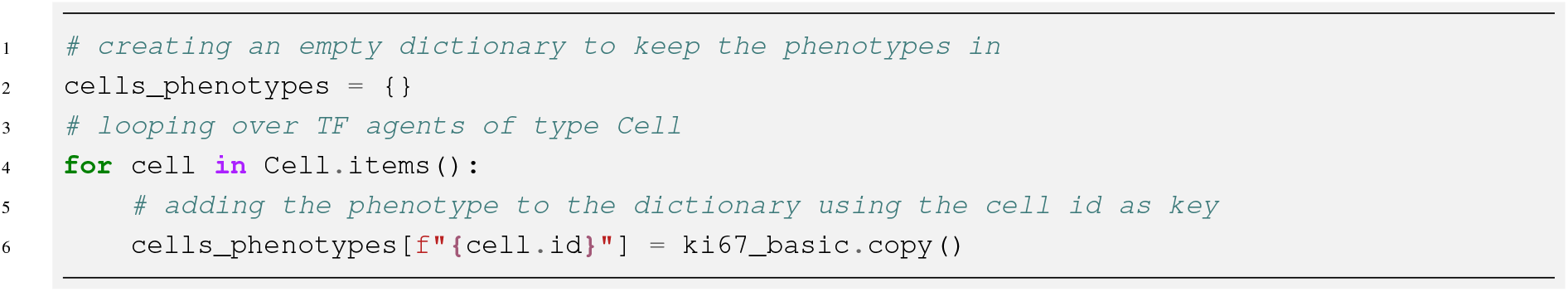
Attaching a Phenotype to a Tissue Forge cell. Listing 2 shows the phenotype initialization.

## 3 Pre-defined Phenotypes

PhenoCellPy comes with several pre-packadged Phenotypes defined. As with the Methods Section (Section 2), we’ve removed most docstrings and comments from code presented in this Section.

### 3.1 Simple Live Cycle

Simplest Phenotype defined, it is a cell-cycle Phenotype of one single Phase. The transition from one Phase to the next (which is the same Phase) is stochastic with an expected duration of *≈* 23*h*, the expected cycle time for a MCF-10A cell line cell [10].

### 3.2 Ki-67 Basic

The Ki-67 Basic is a two Phase cell-cycle phenotype, it is supposed to match experimental data that uses Ki-67, a protein marker for cell proliferation [11]. A cell with a positive Ki-67 marker is in its proliferating state, if there’s no Ki-67 it is in quiescence.

The quiescent Phase (Ki-67 negative) uses an expected Phase duration of 4.59*h*, transition from this Phase is set to be stochastic. The proliferating Phase (Ki-67 positive) uses a fixed duration (*i.e.*, the transition is deterministic) of 15.5*h*. We utilize the same reference values for Phase durations as PhysiCell [4].

### 3.3 Ki-67 Advanced

Ki-67 Advanced adds a post-mitosis Phase to represent the time it takes Ki-67 to degrade post cell division. The proliferating Phase and Ki-67 degradation Phase are both deterministic, with durations of 13*h* and 2.5*h*, respectively. The quiescent (Ki-67 negative) Phase uses the stocastic transition, with an expected duration of 3.62*h*. Again, we utilize the same reference values for Phase durations as PhysiCell [4].

### 3.4 Flow Cytometry Basic

Flow Cytometry Basic is a three Phase live cell cycle. It represents the *G*0*/G*1 *→ S → G*2*/M → G*0*/G*1 cycle. The *G*0*/G*1 phase is more representative of the quiescent phase than the first growth phase, as no growth occurs in this Phase. The Phenotype transitions stochastically from this phase, its expected duration is 5.15*h*. The *S* phase is the phase responsible for doubling the cell volume, transition to the next phase is stochastic, its expected duration is 8*h*. The cell volume growth rate is set to be [total volume growth]/[phase duration]. The G2/M is the pre-mitotic rest phase, the cell divides when exiting this phase. Transition to the next phase is stochastic, its expected duration is 5h. Reference phases durations are from “The Cell: A Molecular Approach. 2nd edition” [12].

### 3.5 Flow Cytometry Advanced

Flow Cytometry Advanced is a four Phase live cell cycle. It represents the *G*0*/G*1 *→ S → G*2 *→ M → G*0*/G*1 cycle. The behaviors The mechanics of the Phases are the same as in Flow Cytometry Basic (Section 3.4) with an added rest Phase (the separation of *G*2 from *M*). All Phase transitions are stochastic, and their expected durations are: 4.98*h*, 8*h*, 4*h*, and 1*h*, respectively. Cell division occurs when exiting Phase *M*.

### 3.6 Apoptosis Standard

Apoptosis Standard is a single Phase dead Phenotype. The cell in this Phenotype sets *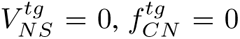*, and *f_F_* = 0 on Phase entry. By doing this, the cell will diminish in volume until it disappears, the rate used for the cytoplasm reduction is *r_CS_* = 1*/*60 *µm*^3^*/min*, for the nucleus it is *r_NS_* = 0.35*/*60 *µm*^3^*/min*, and the fluid change rate is *r_F_* = 3*/*60 *µm*^3^*/min*, volume change rates reference values from [13, 14]. This Phenotype Phase uses a fixed duration of 8.6*h*, the simulated cell should be removed from the simulation domain when the Phase ends.

### 3.7 Necrosis Standard

Necrosis Standard is a two Phase dead Phenotype. The first phase represents the osmotic swell of a necrotic cell, it doesn’t have a set or expected phase duration, instead it uses a custom transition function that monitors the cell volume and transition to the next phase happens when the cell total volume (*V_T_*) reaches its rupture volume (*V_R_*), set to be twice the original volume by default, see Equation 5 and Listing 7. On Phase entry, the solid volumes targets are set to zero (*i.e.*, *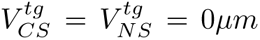*), the target fluid fraction is set to one (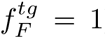), and the target cytoplasm to nuclear ratio is set to zero (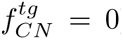). These changes cause the cell to swell. The rates of volume change are: solid cytoplasm *r_CS_* = 3.2*/*60 *×* 10*^−^*^3^ *µm*^3^*/min*, solid nucleus *r_NS_* = 1.3*/*60 *×* 10*^−^*^2^ *µm*^3^*/min*, fluid fraction *r_F_* = 6.7*/*60 *×* 10*^−^*^1^ *µm*^3^*/min*, calcified fraction *r_C_* = 4.2*/*60 *×* 10*^−^*^3^ *µm*^3^*/min*.

The second Phase represents the ruptured cell, it is up to the modeler and modeling framework how the ruptured cell should be represented. For instance, in Tissue Forge [3] the ruptured cell should become several fragment agents, whereas in CompuCell3D [2] the fragmentation can be achieved by setting the ruptured cell contact energy with the medium to be negative. The cell fragments shrink and dissolve into the medium. As a safeguard, this Phase uses a deterministic transition time of 60 days, after which the fragments are flagged for removal. On Phase entry all target volumes are set to zero. The volume change rates are: solid cytoplasm *r_CS_* = 3.2*/*60 *×* 10*^−^*^3^ *µm*^3^*/min*, solid nucleus *r_NS_* = 1.3*/*60 *×* 10*^−^*^2^ *µm*^3^*/min*, fluid fraction *r_F_* = 5*/*60 *×* 10*^−^*^1^ *µm*^3^*/min*, calcified fraction *r_C_* = 4.2*/*60 *×* 10*^−^*^3^ *µm*^3^*/min*.

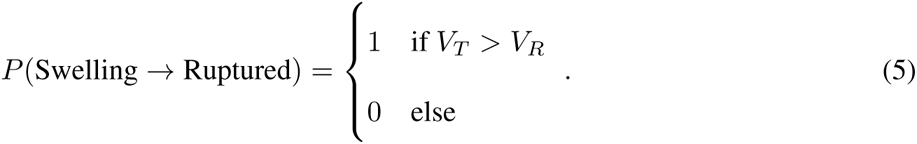

**Listing 7:**
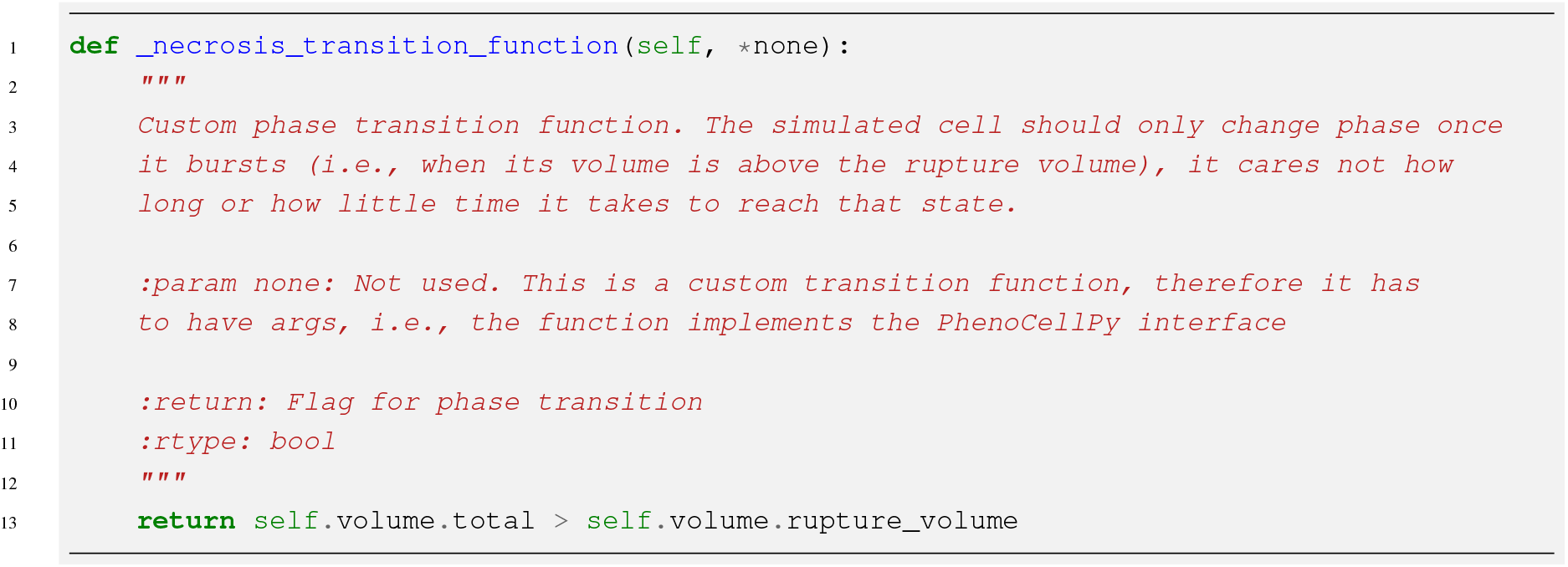
Necrotic Standard’s osmotic swelling Phase transition function.

## 4 Select Examples

### 4.1 CompuCell3D

For CompuCell3D [2] the reccomended method of using PhenoCellPy is to initialize the Phenotype in the steppable start function, and add it as a cell dictionary entry (see Listing 4). Then, in the steppable step function, loop over cells that have a Phenotype model, call the Phenotype time-step, and use the time-step return values as needed (see Listing 1).

See CompuCell3D’s manual [15] for more information on steppables, cell dictionaries, and other CompuCell3D concepts.

#### 4.1.1 Ki-67 Basic Cycle Improved Division

The stock Ki-67 Basic Cycle causes CompuCell3D cells to behave in non-biological ways. This happens because Cellular Potts Model [16] (CompuCell3D’s paradigm) cells are made of several voxels [2], and the proliferating Phase of Ki-67 has a set period. This means that the simulated cells in CompuCell3D may not grow to the doubling volume before they are flagged for cell division. This means that the median cell volume of the cells in the simulation decreases.

Ki-67 Basic Cycle Improved Division fixes this by defining a custom transition function, see Listing 8. It still is a deterministic transition function, however, in addition of monitoring *T* (time spend in phase), it also monitors the simulated cell volume (*V_S_*) and checks if it is bigger than the doubling volume (*V_D_*). The custom transition function evaluates the transition probability as,

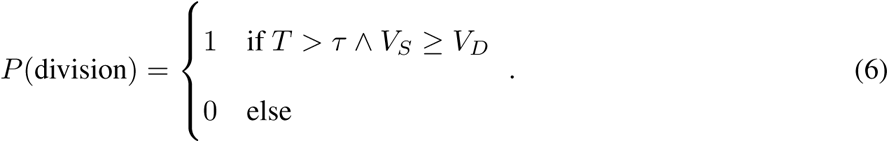

The full implementation of this example is in Supplemental Materials D.

**Listing 8:**
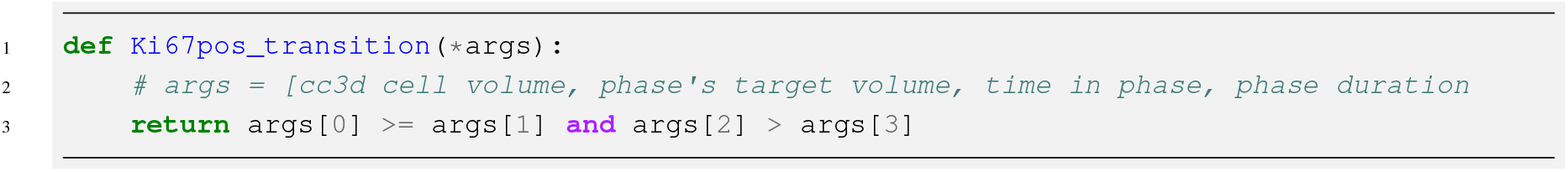
Ki-67 Basic Cycle Improved Division transition function

### 4.2 Tissue Forge

For Tissue Forge [3] the suggested method of using PhenoCellPy is to initialize the Phenotype in a similar way to Tissue Forge’s cell initialization, keep a dictionary of agent IDs as keys and Phenotype models as values, and create a Tissue Forge event [3] to time-step the Phenotype objects. A generic implementation of PhenoCellPy within Tissue Forge can be found in Listing 9. Supplemental Information E shows the implementation of the Ki-67 Basic Cycle Phenotype.

**Listing 9:**
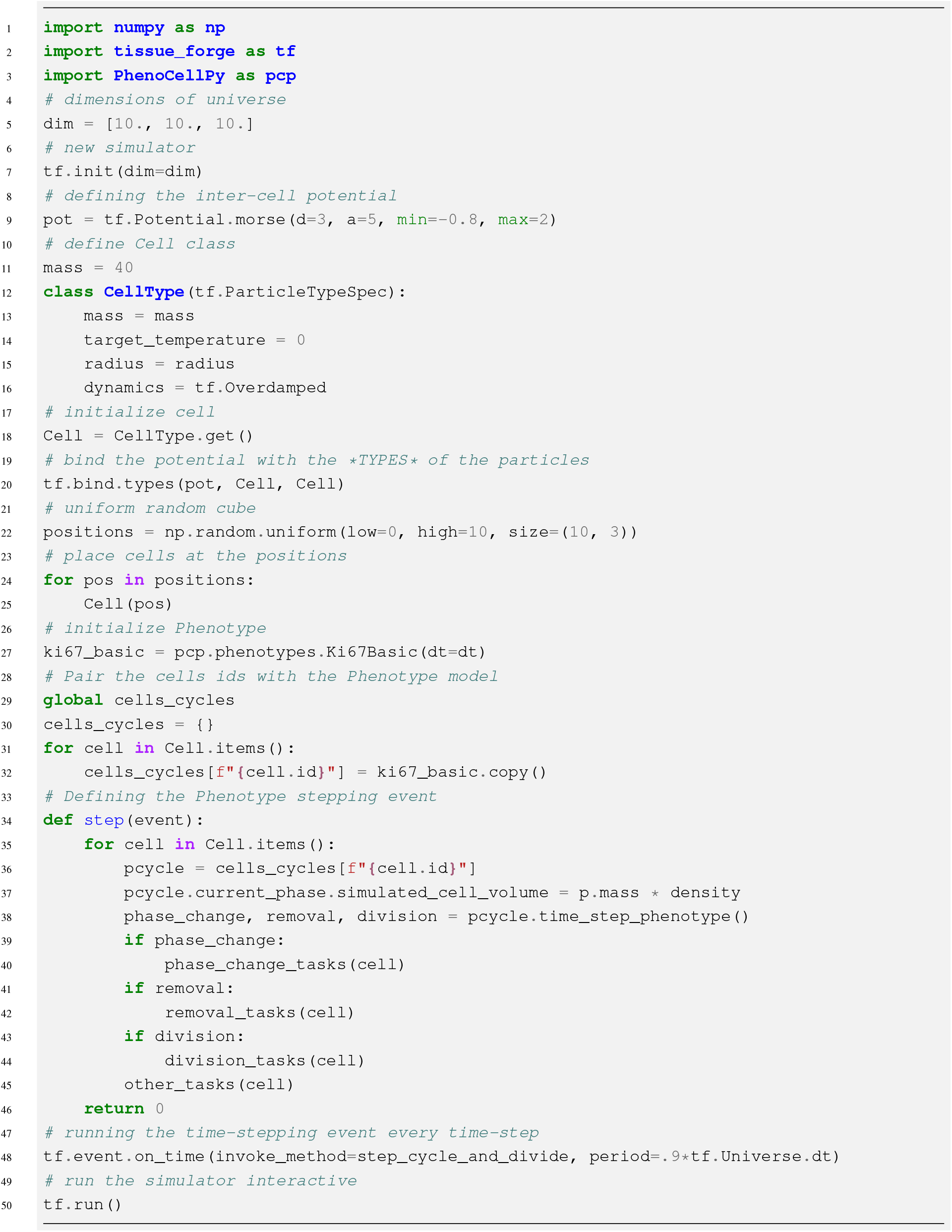
Generic implementation of PhenoCellPy within Tissue Forge

## 5 Selected Results

In this section, we will present selected results from some of the pre-built CompuCell3D and Tissue Forge examples.

### 5.1 PhenoCellPy in CompuCell3D Results

#### 5.1.1 Ki-67 Basic Cycle

We ran the regular Ki-67 Basic cycle (see Section 3.2) and compared it to the Ki-67 Basic cycle improved division (see Section 4.1.1) in CompuCell3D. The simulation here is 2D. In this simulation we start with single cell with a volume of 100 pixels (the pixel to *µm* conversion is set to 24.94*µm/pixel*), therefore they should reach 200 pixels before division. We are using a time-step of 5*min/step*, the usual name for the time-step in CompuCell3D, for historical reasons, is Monte-Carlo Step (MCS). With the regular cycle We see that the median and minimum cell volumes go down in time (Figure 7b), that happens because the mitotic phase transition happens after a set amount of time no matter the cell volume. As cells grow slowly in CompuCell3D they end up being halved before reaching their doubling volume. We see that this effect is worse towards the center of the cell cluster (see Figure 8d), as those cells don’t have room to grow at all. In Figure 8d we can see that many cells are only a few pixels big. We also see that the spatial phase distribution is random in space (see Figure 6e).

**Figure 6:**
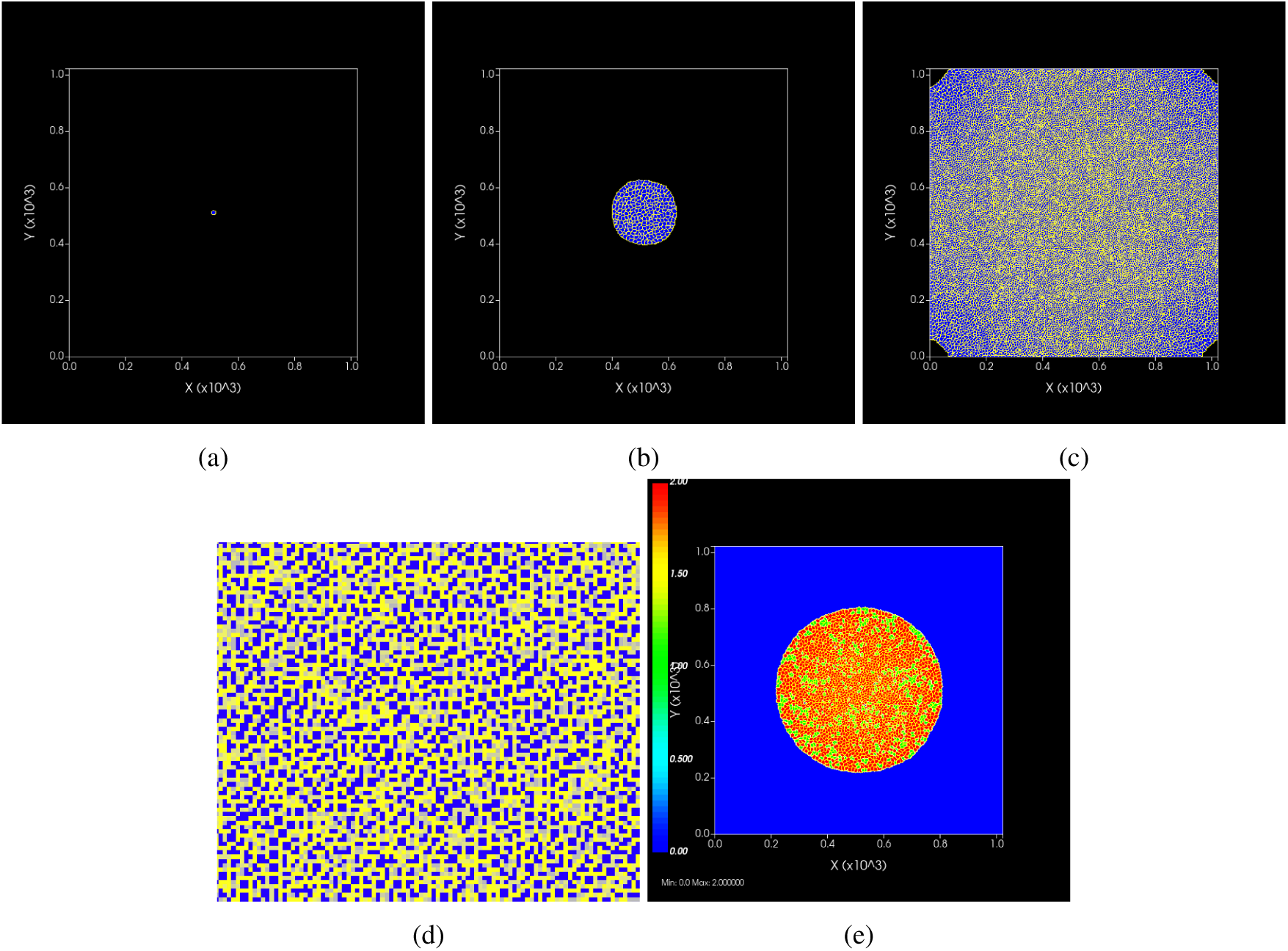
Spatial figures for the CompuCell3D simulation using the regular Ki-67 Basic cycle. a, b, and c) shows the cell cluster with no color overlay. They show the cluster at the start of the simulation, step 2000 and step 4000 respectively. d) Zoom in on the center of image c. e) Color coded cells based on the phase they are in, green for the quiescent phase and red for the proliferating phase. Step 2800.

**Figure 7:**
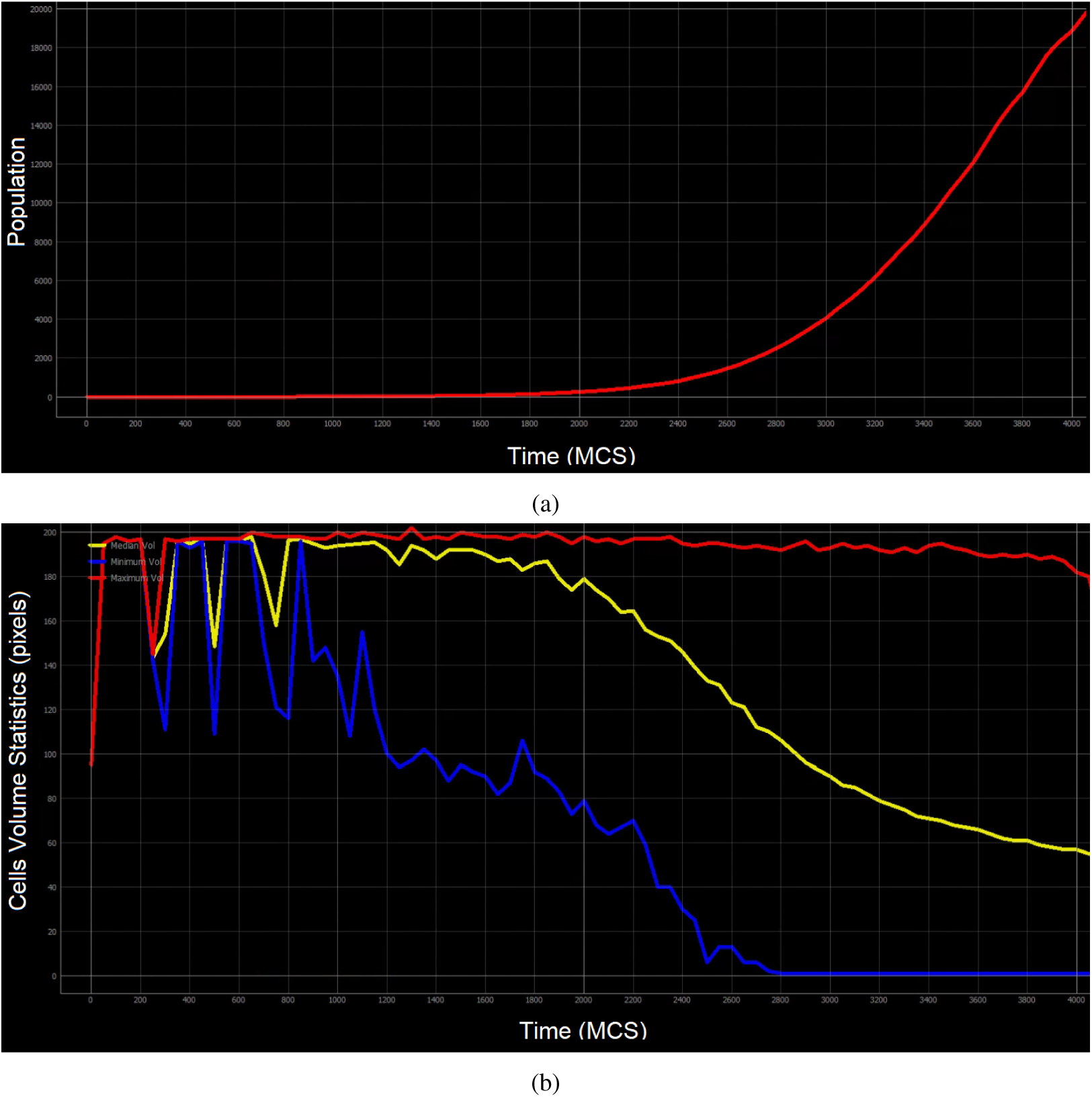
Statistics for the cell population in CompuCell3D using the standard Ki-67 Basic cycle. a) Total cell population. b) Cell population volume statistics. Maximum cell volume in red, median in yellow, and minimum in blue.

**Figure 8:**
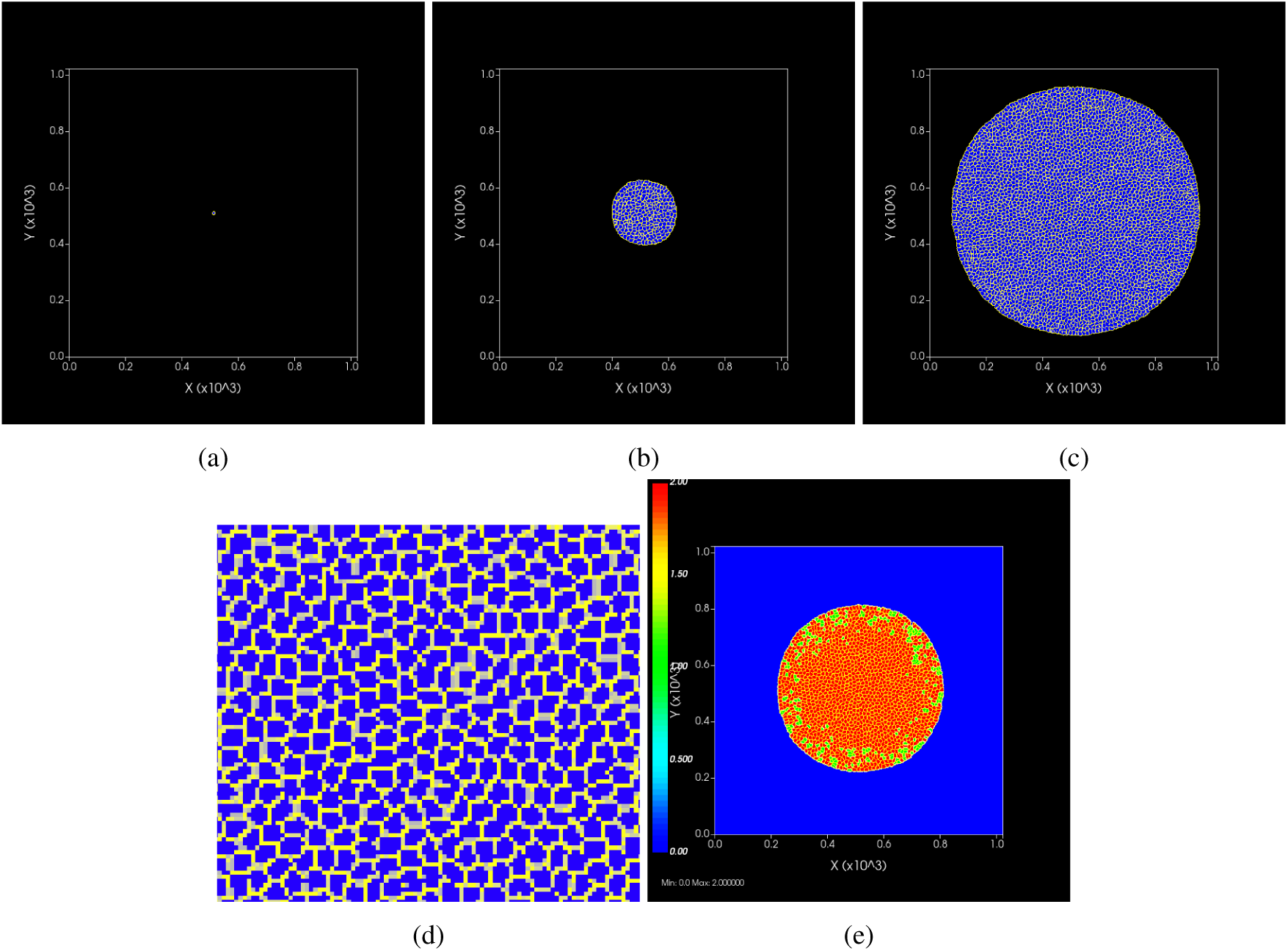
Spatial figures for the CompuCell3D simulation using the regular Ki-67 Basic cycle. a, b, and c) shows the cell cluster with no color overlay. They show the cluster at the start of the simulation, step 2000 and step 4000 respectively. d) Zoom in on the center of image c. e) Color coded cells based on the phase they are in, green for the quiescent phase and red for the proliferating phase. Step 3200.

In contrast, when using the modified transition we account for the simulated cell volume. Now the reduction in the median volume is small and happens due to overcrowding and cells pushing each other, and that the total population growth was much smaller (3800 cells at step 4000, see Figure 9a, versus 19000, see Figure 7a). We do not see overly small cells (see Figure 8d). We also see that cells towards the center of the cell cluster stay in the proliferating phase (see Figure 8e), unable to reach their doubling volume and divide.

**Figure 9:**
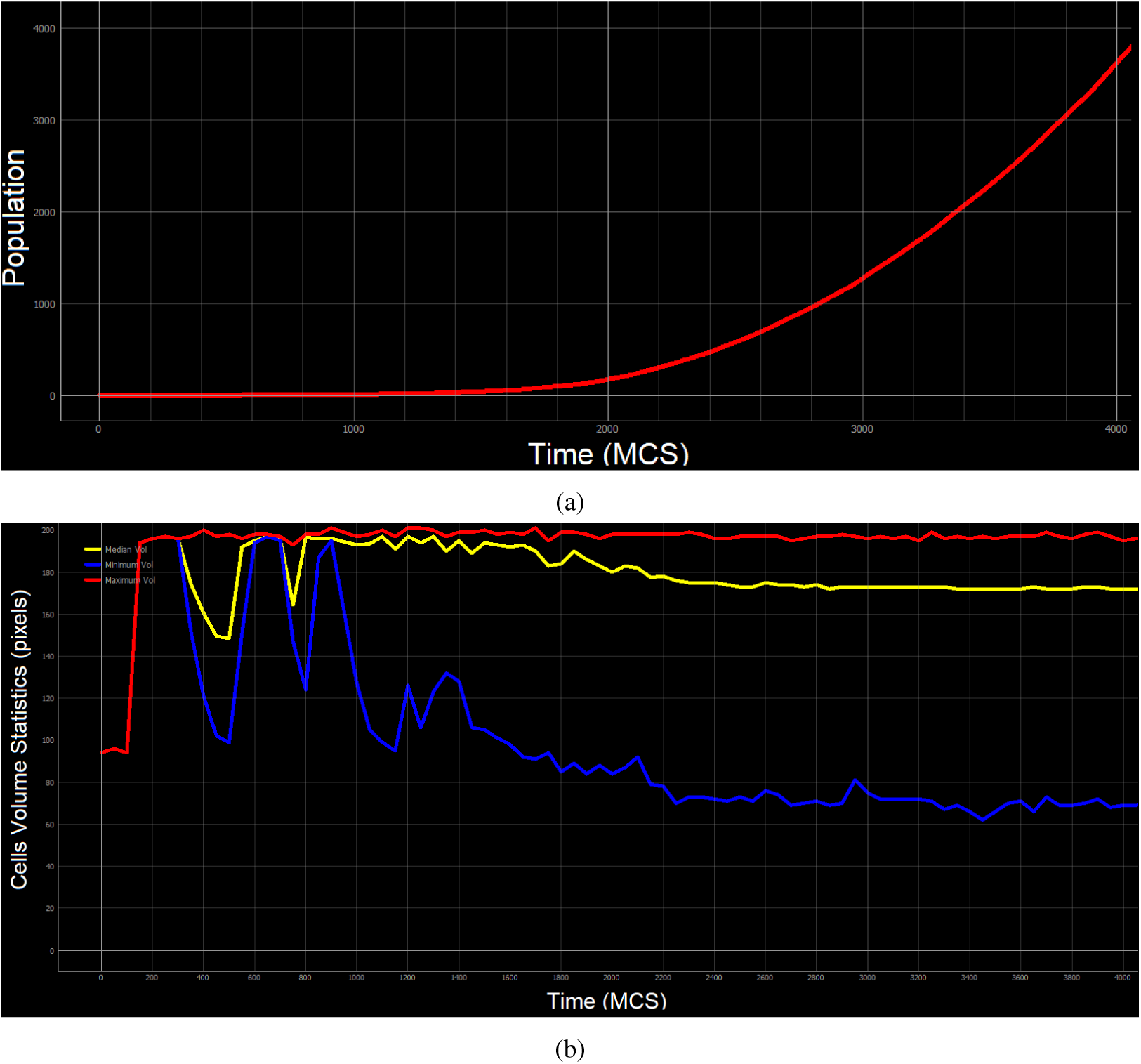
Statistics for the cell population in CompuCell3D using the modified Ki-67 Basic cycle. a) Total cell population. b) Cell population volume statistics. Maximum cell volume in red, median in yellow, and minimum in blue.

#### 5.1.2 Necrosis standard

We ran the necrotic phenotype in a 2D CompuCell3D simulation, we simulate 170 cells plated in a petri dish. We select ten of those cells to undergo necrosis. We keep the initial volume of the cells 100 pixels (with the same pixel to *µm* relation) and the same time-step duration of 5*min/step*. The necrotic cells increase in volume linearly (see Figure 11) during the hydropic (osmotic) swell phase until they reach their bursting volume. After bursting (see Figure 10) their volume decreases rapidly at first, going near zero almost immediately, and then keeps decreasing more slowly (see Figure 11). This happens because the PhenoCellPy model sets the target volume of the cell to 0 when entering the ruptured phase, however, CompuCell3D works by minimizing the energy of the system, and the cell fragments have a negative contact energy with the medium (see CompuCell3D’s [2] manuals for a definition of the contact energy).

**Figure 10:**
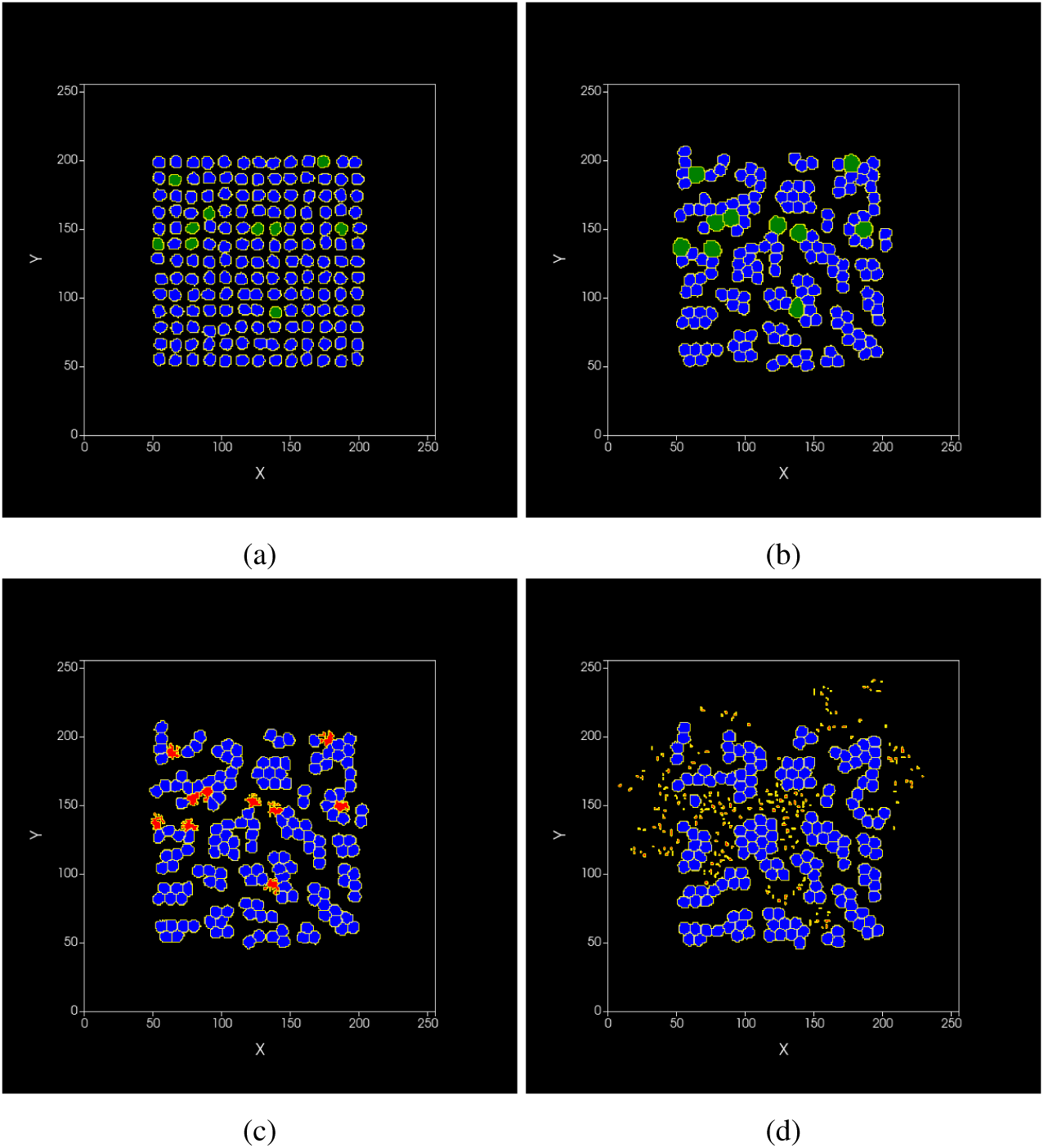
Snapshots of the CompuCell3D simulation for the necrotic phenotype, necrotic cells in green, healthy cells in blue, fragmented cells in red. a) simulation start. b) Necrotic cells at maximum volume. c) Moment the necrotic cells bursts. d) end of the simulation.

**Figure 11:**
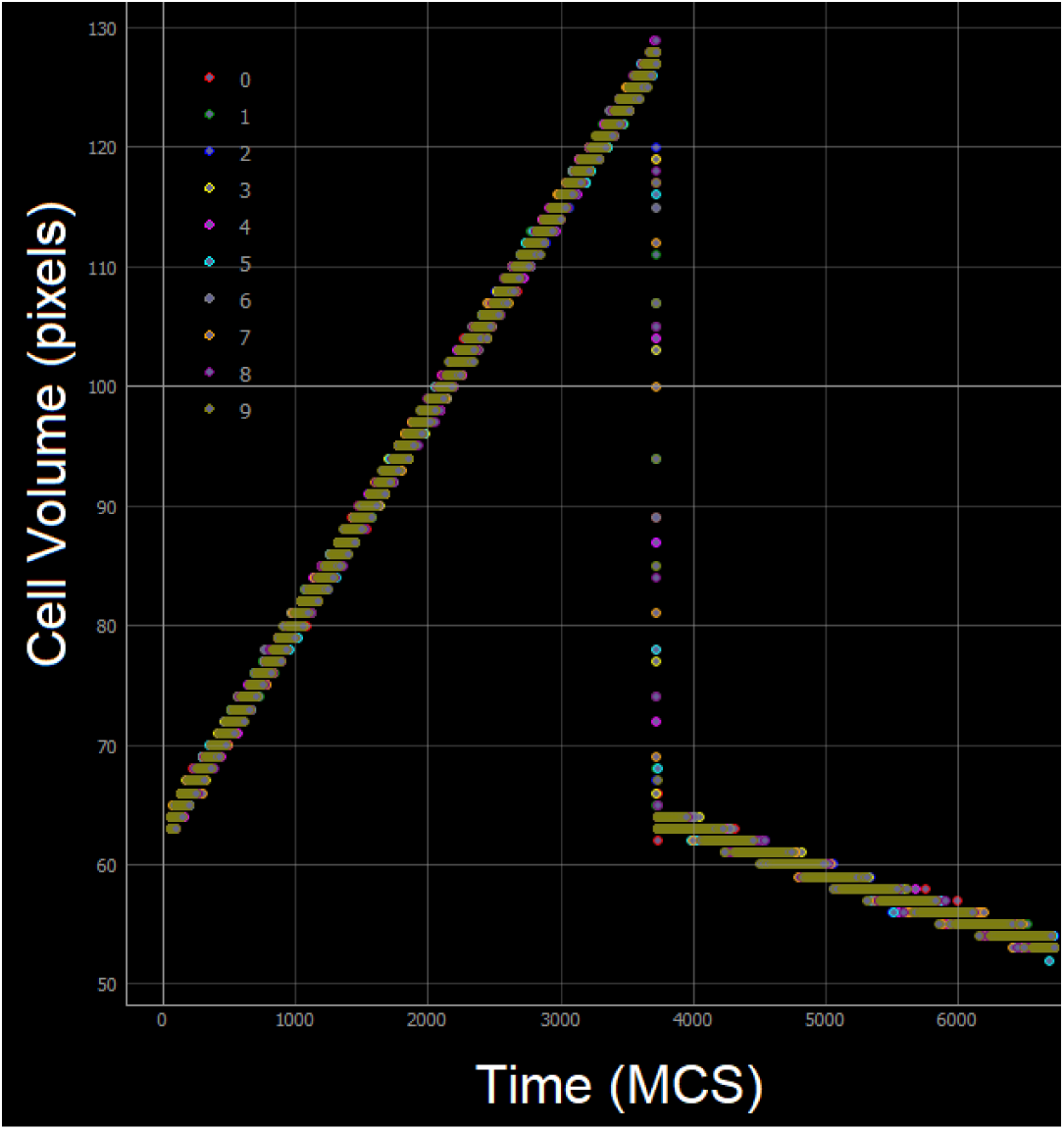
Necrotic cells volumes evolution in time. Each necrotic cell volume is plotted individuality

### 5.2 Tissue Forge

#### 5.2.1 Ki-67 Basic Cycle

For the Tissue Forge Ki-67 Basic Cycle simulation we used a time-step duration of 10 min/step, as Tissue Forge is off-lattice we don’t need a space conversion factor. This simulation is 3D, and we see that the cell cluster at the center (see Figure 12).

**Figure 12:**
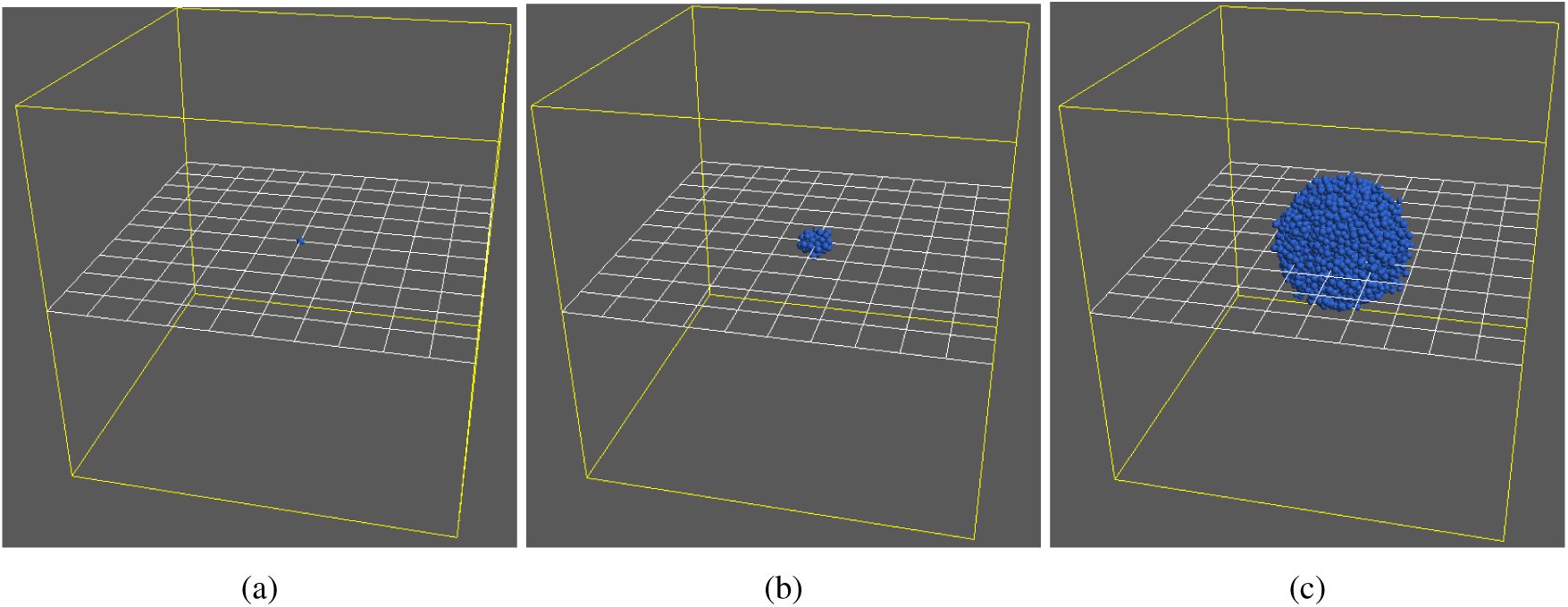
Cells space configuration in the Tissue Forge Ki-67 Basic cycle model. a) Simulation start. b) Day 7. c) Day 15 (simulation end).

Unlike in CompuCell3D, cells in Tissue Forge are soft spheres and can reach their desired volume almost instantly. Therefore, we don’t need the modification to the division transition. The median and minimum cell volume stay steady (see Figure 13b). The median cell volume stays close to the maximum cell volume (see Figure 13b) because the cells double in volume very fast in Tissue Forge, the cells then stay at their doubling volume until the phase duration has transpired. As before, the population growth is exponential (see Figure 13a).

**Figure 13:**
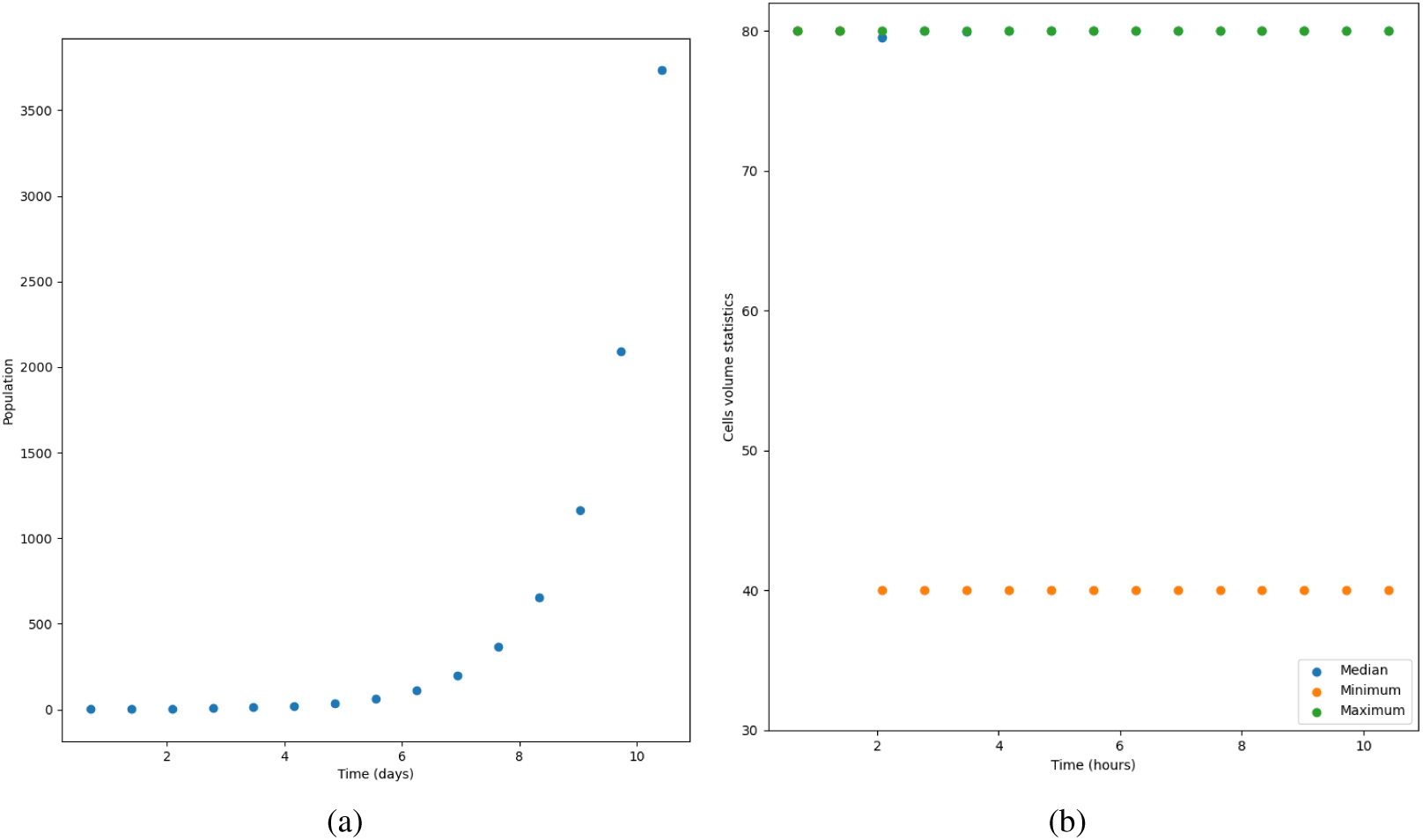
Cell population statistics for the Tissue Forge model using PhenoCellPy’s Ki-67 Basic Cycle phenotype. a) Total cell population. b) Cells’ volume statistic.

## 6 Discussion

The PhenoCellPy’s embedded modeling package allows modelers to easily create sequences of cell behaviors and attach them to agent in agent-based models. PhenoCellPy is accessible, intuitive and enables modelers to add complexity to their model without much overhead. It implements several biological concepts, such as how to switch from one behavior to the next, how the different volumes of the cell should be modeler (*e.g.,* cytoplasmic and nuclear), and makes sure time-scales are respected. PhenoCellPy also makes the creation of new behaviors and sequences of behaviors easy.

PhenoCellPy is open-source and freely available under the BSD 3-Clause License (https://github.com/JulianoGianlupi/PhenoCellPy/blob/main/LICENSE).

### 6.1 Installation

PhenoCellPy’s *alpha* version does not have any installer. To use it you should clone or download its GitHub repository [5] and add its folder to the simulation’s system path. E.g., see Listing 10.

**Listing 10:**
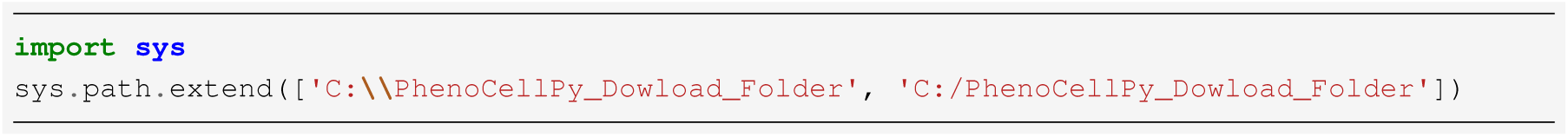
How to add PhenoCellPy to the simulation’s system path.

### 6.2 Planned features

We currently have these planned features:

- Interface (API) classes for CompuCell3D and Tissue Forge
- Template interface classes
- Automatic inter-cell heterogeneity [17]
- Randomization of the initial Phase of the Phenotype
- Conda or pip distribution
- API for environment interaction (*e.g.*, detection of oxygen levels in the environment, detection of neigboring cells)
- ”Super-Phenotypes,” phenotypes made from more than one Phenotype class, and methods for switching between them

### 6.3 Requirements

PhenoCellPy only requirements, besides Python 3 support, are that NumPy and SciPy be available inside the modeling framework PhenoCellPyis embedded in.

## Footnotes

1 Nomenclature in biology is diverse, with several different definitions for the same term. We define what phenotype and phase mean in the context of PhysiCell and PhenoCellPy in our text.

2 see BioPortal’s ontology definition for hydropic

3 see BioPortal’s ontology definition for lysed

