## Supplemental Materials for "PhenoCellPy: A Python package for biological cell behavior modeling"

### A PhenoCellPy Python implementation

Besides defining several methods and functions to drive PhenoCellPy, we allow users to define custom functions. The user-definable functions are: `entry_function(*args)`, `exit_function(*args)`, `arrest_function(*args)`, `user_phenotype_time_step(*args)`, `user_phase_time_step(*args)`. Which are executed at Phase entry, Phase exit, to determine if a cell should exit the Phenotype and enter senescence (see Sections 2.2 and 2.3), and at each time-step. We also allow the user to define the Phase transition function (see Section 2.2.1). All functions that can be user-defined in PhenoCellPy **must** accept any number of optional parameters (*i.e.*, be a ”python args” function). For instance, for viral infection the end of the eclipse phase will occur when a threshold of internal viral load is reached [1], so the modeler would pass the intra-cellular viral load and the threshold as arguments to the function. Some pre-packaged Phases (*e.g.*, `NecrosisSwell`, `S`) use the custom entry function to change their Cell Volume model target volumes.

We would like to note that, for formatting reasons, we've removed the docstrings and most comments from the functions presented in this section. The actual source-code for PhenoCellPy contains docstrings for all functions and comments where necessary.

#### A.1 Cellular Phase

The Phase is the ”base unit” of the Phenotype. Each Phase has as attributes a descriptive name (*e.g.*, S, G, M, necrotic swelling), an index (*i.e.* in which position of the sequence of Phases it is), the index of the previous and next Phases, the time-step length ($dt$), the name of the time unit (*e.g.,* second, minute), how long the Phase should be ($\tau$, ”`phase_duration`” in the code, see Section 2.2.1 and A.1.4), a flag setting the transition to the next Phase to be stochastic or deterministic (see Section 2.2.1 and A.1.4), a flag for mitosis or meiosis at phase exit, a flag for removal from the simulation at phase exit (*i.e.*, the cell dies, is killed, leaves the simulated domain). The Phase class also keeps track of the amount of time the cell has spent in the current Phase ($T$), and (optionally) the volume of the cell in the simulation. Although the cell volume definition and update is handled by the Cell Volume class (Section 2.1), the volume change rates is an attribute of the Phase class. It's important to note that the volume of the cell defined by PhenoCellPy's volume class may differ from the volume of the cell in the simulation.

The Phase class also has several functions defined, to time-step the model (Section A.1.2), to update the cellular volume model (Section 2.1), to double the target volume of the cellular volume model, and

to evaluate if the transition to the next Phase should occur (Section 2.2.1 and A.1.4). It also has placeholder functions that can be replaced by user-defined ones, one that should be executed upon Phase entry (`entry_function(*args)`), one just before Phase exit (`exit_function(*args)`), one for exiting the Phenotype and entering senescence (`arrest_function(*args)`), and one that is executed at each time-step
(`user_phase_time_step(*args)`), see Section A.2.4.

##### A.1.1 Phase class initialization

The Phase `__init__` function performs some checks on attribute values, *e.g.* $dt > 0$, a negative or zero time-step makes no sense, the custom functions should be functions, and so on. Initializes attributes to set values, and initializes the Cellular Volume model class. The Phase `__init__` function is in Supplemental Materials B.

##### A.1.2 Phase time-step

The Phase's time-step function is responsible for incrementing $T$ (total time the cell's spent in the current Phase, `time_in_phase` in the code) by $dt$. Then it calls the volume update function and checks if the cell exits the Phenotype and goes into senescence. Finally, it calls the Phase transition function (the default deterministic or stochastic, or a user-defined function). It returns a tuple of two boolean flags, the first flags if the cell should go to the next Phase in the Phenotype, the second if the cell should exit the Phenotype and enter senescence. The Phase time-step function is shown in Listing 11.

##### A.1.3 Phases volume update

The volume update itself is handled by the Cell Volume class. As the volume change rates are an attribute of the Phase class, however, the Phase class has its own intermediary update volume function. This intermediary function's job is to pass the rates to the Cell Volume's update volume function. The intermediary function is shown in Listing 12.

##### A.1.4 Phase Transition

The transition from one phase to the next can be either deterministic (with a set period) or stochastic (with a set transition rate) by setting the Phase's class attribute "`fixed_duration`" to be True (deterministic) or False (stochastic). Based on the flag, the Phase class sets its transition check function to be either `_transition_to_next_phase_deterministic` (for the deterministic case), Listing 13, or

```python
def time_step_phenotype(self):
        self.time_in_phase += self.dt
        self.update_volume()
        transition_to_index = None
        if self.user_phase_time_step is not None:
            self.user_phase_time_step(*self.user_phase_time_step_args)
        if self.arrest_function is not None:
            exit_phenotype = self.arrest_function(*self.exit_function_args)
            go_to_next_phase_in_phenotype = False
            return go_to_next_phase_in_phenotype, exit_phenotype, transition_to_index
        else:
            exit_phenotype = False
        go_to_next_phase_in_phenotype = self.check_transition_to_next_phase_function(
            *self.check_transition_to_next_phase_function_args)
        if hasattr(go_to_next_phase_in_phenotype, "len") and \
            len(go_to_next_phase_in_phenotype) > 1:
                transition_to_index = go_to_next_phase_in_phenotype[1]
                go_to_next_phase_in_phenotype = go_to_next_phase_in_phenotype[0]
        if go_to_next_phase_in_phenotype and self.exit_function is not None:
            self.exit_function(*self.exit_function_args)
            return go_to_next_phase_in_phenotype, exit_phenotype, transition_to_index
        return go_to_next_phase_in_phenotype, exit_phenotype, transition_to_index
```

Listing 11: Phase class time-step function.

```python
def update_volume(self):
    self.volume.update_volume(self.dt, self.fluid_change_rate,
        self.nuclear_volume_change_rate, self.cytoplasm_volume_change_rate,
        self.calcification_rate)
```

Listing 12: Phase class volume update intermediary function.

`_transition_to_next_phase_stochastic` (for the stochastic case), Listing 14. By default, `fixed_duration` is `False`, *i.e.*, the default behavior is to use the stochastic transition. A user can also define their own transition function that can take into account other factors. As the Phase transition function can be defined by the user, our default transition functions must also have `*args` as its argument.

```python
def _transition_to_next_phase_deterministic(self, *none):
    return self.time_in_phase > self.phase_duration
```

Listing 13: Deterministic transition function

```python
def _transition_to_next_phase_stochastic(self, *none):
    prob = 1 - exp(-self.dt / self.phase_duration)
    return uniform() < prob
```

Listing 14: Stochastic transition function

#### A.2 Cellular Phenotype

The Phenotype class has as attributes a descriptive name (*e.g.*, Standard necrosis model, Flow Cytometry Basic), the time-step length ($dt$), the name of the time unit (*e.g.,* second, minute), a list of Phases that make the Phenotype, the index of the Starting Phase, an optional "stand-alone" senescent Phase (i.e., a Phase that is outside the Phenotype cycle), the current Phase, and the amount of time spent in this Phenotype. Note that some of these attributes are shared with the Phase class, the Phenotype class should pass these shared attributes to its component Phases upon initialization.

This class has methods to time-step the phenotype model (Section A.2.2), to perform user-defined time-step tasks, to change the phenotype phase to an arbitrary phase of the phenotype cycle (Section A.2.3), to go to the next phase in the cycle (Section A.2.3), and to go to a non-changing senescent phase (Section A.2.4).

##### A.2.1 Phenotype class initialization

The Phenotype `__init__` function performs sanity checks and initializes attributes. Initialization of component Phases should be made in the Phenotype `__init__` function. The Phenotype class `__init__` function is shown in Listing 15.

##### A.2.2 Phenotype time-step

The first thing the Phenotype time-step function (Listing 16) does is check if the Phenotype has just started (i.e., the time spent thus far in the Phenotype is 0), and if so it calls the initial Phase entry function. Then it increments the "amount of time spent in this Phenotype" attribute ( `time_in_cycle` in the code) by $dt$, calls the current Phase's time-step function (Section A.1.2, and checks if the Phenotype should move to the next Phase or go to quiescence (as determined by the Phase's time-step).

If the Phenotype should go to the next Phase it calls the `go_to_next_phase` function (Section A.2.3), if it should go to quiescence it calls the `go_to_quiescence` function (Section A.2.4). The time-step function returns a tuple of three boolean flags, the values of which can be determined by the `go_to_next_phase` function. The first flag of the tuple says if the Phenotype has changed Phases, the second if the simulated cell should be removed from the simulation, and the third if the simulated cell has undergone cell division.

```python
def __init__(self, name: str = "unnamed", dt: float = 1, time_unit: str = "min",
    phases: list = None, senescent_phase: Phases.Phase or False = None,
    starting_phase_index: int = 0):
    self.name = name
    self.time_unit = time_unit
    if dt <= 0 or dt is None:
        raise ValueError(f"'dt' must be greater than 0. Got {dt}.")
    self.dt = dt
    if phases is None:
        self.phases = [Phases.Phase(previous_phase_index=0,
            next_phase_index=0, dt=self.dt, time_unit=time_unit)]
    else:
        self.phases = phases
    if senescent_phase is None:
        self.senescent_phase = Phases.SenescentPhase(dt=self.dt)
    elif senescent_phase is not None and not senescent_phase:
        self.senescent_phase = False
    elif not isinstance(senescent_phase, Phases.Phase):
        raise ValueError(
            f"`senescent_phase` must Phases.Phase object, False, or None."
            f" Got {senescent_phase}")
    else:
        self.senescent_phase = senescent_phase
    if starting_phase_index is None:
        starting_phase_index = 0
    self.current_phase = self.phases[starting_phase_index]
    self.time_in_phenotype = 0
```

Listing 15: Phenotype class `__init__` function

##### A.2.3 Switching Phases

The `set_phase` function (Listing 17) is responsible for switching the Phase of a Phenotype, it does so by setting the `current_phase` to be the phase of index $i$. It also ensures that the Cell Volume sub-class attributes are correct in the case the cell has changed volume while in the current Phase and resets T (the time spent in the Phase) to zero. Finally, it calls the new Phase entry function if there is one.

```
def time_step_phenotype(self):
            if not self.time_in_phenotype and \
                self.current_phase.entry_function is not None:
                    self.current_phase.entry_function(
                        *self.current_phase.entry_function_args)
        if self.user_phenotype_time_step is not None:
            self.user_phenotype_time_step(
                *self.user_pheno_time_step_args)
        self.time_in_phenotype += self.dt
        go_next_phase, exit_phenotype, transition_to_index = \
            self.current_phase.time_step_phase()
        if go_next_phase:
            if transition_to_index is not None:
                phase_idx = self.current_phase.index
                old_next_phase_idx = self.current_phase.next_phase_index
                self.current_phase.next_phase_index = transition_to_index
            else:
                phase_idx = None
            changed_phases, cell_removed, cell_divides = self.go_to_next_phase()
            if phase_idx is not None:
                self.phases[phase_idx].next_phase_index  = old_next_phase_idx
            return changed_phases, cell_removed, cell_divides
        elif exit_phenotype:
            self.go_to_senescence()
            changed_phases, cell_removed, cell_divides = (True, False, False)
            return changed_phases, cell_removed, cell_divides
        changed_phases, cell_removed, cell_divides = (False, False, False)
        return changed_phases, cell_removed, cell_divides
```

Listing 16: Phenotype time-step function

```
def set_phase(self, idx):
    # get the current cytoplasm, nuclear, calcified volumes
    cyto_solid = self.current_phase.volume.cytoplasm_solid
    cyto_fluid = self.current_phase.volume.cytoplasm_fluid
    nucl_solid = self.current_phase.volume.nuclear_solid
    nucl_fluid = self.current_phase.volume.nuclear_fluid
    calc_frac = self.current_phase.volume.calcified_fraction
    # get the target volumes
    target_cytoplasm_solid = self.current_phase.volume.cytoplasm_solid_target
    target_nuclear_solid = self.current_phase.volume.nuclear_solid_target
    target_fluid_fraction = self.current_phase.volume.target_fluid_fraction
    # set parameters of next phase
    self.phases[idx].volume.cytoplasm_solid = cyto_solid
    self.phases[idx].volume.cytoplasm_fluid = cyto_fluid
    self.phases[idx].volume.nuclear_solid = nucl_solid
    self.phases[idx].volume.nuclear_fluid = nucl_fluid
    self.phases[idx].volume.calcified_fraction = calc_frac
    self.phases[idx].volume.cytoplasm_solid_target = target_cytoplasm_solid
    self.phases[idx].volume.nuclear_solid_target = target_nuclear_solid
    self.phases[idx].volume.target_fluid_fraction = target_fluid_fraction
    # set phase
    self.current_phase = self.phases[idx]
    self.current_phase.time_in_phase = 0
    if self.current_phase.entry_function is not None:
        self.current_phase.entry_function(*self.current_phase.entry_function_args)
```



Listing 17: Phenotype Class `set_phase` function.

The `go_to_next_phase` (Listing 18) switches to the next Phase by calling `set_phase` with the current Phase's "next phase index" attribute. Before switching Phases, it fetches the boolean flags for division and removal (*e.g.*, death, leaving the simulation domain) at Phase exit. It returns a tuple of three boolean flags: Phase change, cell removal from the simulation, and cell division.

```python
def go_to_next_phase(self):
    changed_phases = True
    divides = self.current_phase.division_at_phase_exit
    removal = self.current_phase.removal_at_phase_exit
    self.set_phase(self.current_phase.next_phase_index)
    return changed_phases, removal, divides
```

Listing 18: Phenotype Class `go_to_next_phase` function

##### A.2.4 Leaving the Phenotype

To exit the Phenotype the modeler has to define the optional arrest function which is a member of the Phase class (Section 2.2). The arrest function is called by the Phase time-step function (Section A.1.2) and its return value is used by the Phenotype time-step function to exit the Phenotype cycle.

If the arrest function returns `True`, the Phenotype time-step calls `go_to_senescence`. `go_to_senescence` checks that the Phenotype attribute `senescent_phase` is a class of type Phase, if that's the case it sets the current phase to be the special senescent Phase. If that's not the case the function will simply return early.

Before the Phase change, `go_to_quiescence` saves the volume parameters to temporary variables to keep them as they are after the change. As this is a change to a senescent Phase, all target volumes of the Cell Volume sub-class are set to be the current actual volumes (see Section 2.1 for definition of target volume and actual volumes). It resets `time_in_phase` to be 0. The `go_to_quiescence` function is shown in Listing 19.

```python
def go_to_senescence(self):
    if not isinstance(self.senescent_phase, Phases.Phase):
        return
    # get the current cytoplasm, nuclear, calcified volumes
    cyto_solid = self.current_phase.volume.cytoplasm_solid
    cyto_fluid = self.current_phase.volume.cytoplasm_fluid
    nucl_solid = self.current_phase.volume.nuclear_solid
    nucl_fluid = self.current_phase.volume.nuclear_fluid
    calc_frac = self.current_phase.volume.calcified_fraction
    # setting the senescent phase volume parameters.
    # As the cell is now senescent it shouldn't want to change its
    # volume, so we set the targets to be the current measurements
    self.senescent_phase.volume.cytoplasm_solid = cyto_solid
    self.senescent_phase.volume.cytoplasm_fluid = cyto_fluid
    self.senescent_phase.volume.nuclear_solid = nucl_solid
    self.senescent_phase.volume.nuclear_fluid = nucl_fluid
    self.senescent_phase.volume.nuclear_solid_target = nucl_solid
    self.senescent_phase.volume.cytoplasm_solid_target = cyto_solid
    self.senescent_phase.volume.calcified_fraction = calc_frac
    self.senescent_phase.volume.target_fluid_fraction = (cyto_fluid + nucl_fluid) / \
                        (nucl_solid + nucl_fluid + cyto_fluid + cyto_solid)
    # set the phase to be senescent
    self.current_phase = self.senescent_phase
    self.current_phase.time_in_phase = 0
```

Listing 19: Phenotype Class `go_to_quiescence` function.

#### A.3 Cellular Volume

The update volume function is shown in Listing 20. It takes as arguments, in order, the time-step length ($dt$), $r_F$, $r_{NS}$, $r_{CS}$, and $r_C$. For stability reasons, we use SciPy's `odeint` function [18] to solve the volume dynamics, together with an intermediary function `volume_relaxation` (see Listing 21).

```python
def update_volume(self, dt, fluid_change_rate, nuclear_volume_change_rate,
    cytoplasm_volume_change_rate, calcification_rate):
        dt_array = array([0, dt])
        self.fluid = odeint(self.volume_relaxation, self.fluid, dt_array,
                        args=(fluid_change_rate,
                            self.target_fluid_fraction * self.total))[-1][0]
        self.nuclear_fluid = (self.nuclear / (self.total + 1e-12)) * self.fluid
        self.cytoplasm_fluid = self.fluid - self.nuclear_fluid
        self.nuclear_solid = odeint(self.volume_relaxation, self.nuclear_solid, dt_array,
                            args=(nuclear_volume_change_rate,
                                  self.nuclear_solid_target))[-1][0]
        self.cytoplasm_solid_target = self.target_cytoplasm_to_nuclear_ratio * \
                                  self.nuclear_solid_target
        self.cytoplasm_solid = odeint(self.volume_relaxation,
                                  self.cytoplasm_solid, dt_array,
                                  args=(cytoplasm_volume_change_rate,
                                        self.cytoplasm_solid_target))[-1][0]
        self.solid = self.nuclear_solid + self.cytoplasm_solid
        self.nuclear = self.nuclear_solid + self.nuclear_fluid
        self.cytoplasm = self.cytoplasm_fluid + self.cytoplasm_solid
        self.calcified_fraction = odeint(self.volume_relaxation,
                                     self.calcified_fraction, dt_array,
                                     args=(calcification_rate, 1))[-1][0]
        self.total = self.cytoplasm + self.nuclear
        self.fluid_fraction = self.fluid / (self.total + 1e-12)
```

Listing 20: Cell Volume class `update_volume` function.

```python
@staticmethod
def volume_relaxation(current_volume, t, rate, target_volume):
    dvdt = rate * (target_volume - current_volume)
    return dvdt
```

Listing 21: Intermediary `volume_relaxation` function.

##### A.3.1 Cellular Volume defaults and init function

The Cellular Volume `__init__` function checks if the user defined custom values, checks that they are sensible (e.g., that there are no negative volume, or fractional volumes $\notin [0, 1]$), and initializes the class' attributes.

#### B Phase init function

```python
def __init__(self, index: int = None, previous_phase_index: int = None,
    next_phase_index: int = None, dt: float = None, time_unit: str = "min",
    name: str = None, division_at_phase_exit: bool = False,
    removal_at_phase_exit: bool = False, fixed_duration: bool = False,
    phase_duration: float = 10, entry_function=None, entry_function_args: list = None,
    exit_function=None, exit_function_args: list = None, arrest_function=None,
    arrest_function_args: list = None, transition_to_next_phase=None,
    transition_to_next_phase_args: list = None,
    simulated_cell_volume: float = None, cytoplasm_volume_change_rate=None,
    nuclear_volume_change_rate=None, calcification_rate=None,
    target_fluid_fraction=None, nuclear_fluid=None, nuclear_solid=None,
    nuclear_solid_target=None, cytoplasm_fluid=None, cytoplasm_solid=None,
    cytoplasm_solid_target=None, target_cytoplasm_to_nuclear_ratio=None,
    calcified_fraction=None, fluid_change_rate=None, relative_rupture_volume=None):

    if index is None:
        self.index = 0
    else:
        self.index = index
    self.previous_phase_index = previous_phase_index
    self.next_phase_index = next_phase_index
    self.time_unit = time_unit
    if dt is None or dt <= 0:
        raise ValueError(f"'dt' must be greater than 0. Got {dt}.")
    self.dt = dt
    if name is None:
        self.name = "unnamed"
    else:
        self.name = name
    self.division_at_phase_exit = division_at_phase_exit
    self.removal_at_phase_exit = removal_at_phase_exit
    self.fixed_duration = fixed_duration
    if phase_duration <= 0:
        raise ValueError(f"'phase_duration' must be greater than 0. Got {phase_duration}")
    self.phase_duration = phase_duration
    self.time_in_phase = 0
    self.entry_function = entry_function
    self.entry_function_args = entry_function_args
```

```python
        self.exit_function = exit_function
        self.exit_function_args = exit_function_args
        if self.exit_function is not None and not (type(self.exit_function_args) == list or
                                                   type(self.exit_function_args) == tuple):
            raise TypeError("Exit function defined but no args given. Was expecting "
                            f"'exit_function_args' to be a list or tupple, got {type(
                                                                         exit_function_args)}.
                                                                         ")

        self.arrest_function = arrest_function
        self.arrest_function_args = arrest_function_args
        if self.arrest_function is not None and type(self.arrest_function_args) != list:
            raise TypeError("Arrest function defined but no args given. Was expecting "
                            f"'arrest_function_args' to be a list, got {type(
                                                                         arrest_function_args)
                                                                         }.")

        if transition_to_next_phase is None:
            self.transition_to_next_phase_args = [None]
            if fixed_duration:
                self.transition_to_next_phase = self._transition_to_next_phase_deterministic
            else:
                self.transition_to_next_phase = self._transition_to_next_phase_stochastic
        else:
            if type(transition_to_next_phase_args) != list:
                raise TypeError("Custom exit function selected but no args given. Was
                                                             expecting "
                                "'transition_to_next_phase_args' to be a list, got "
                                f"{type(transition_to_next_phase_args)}.")
            self.transition_to_next_phase_args = transition_to_next_phase_args
            self.transition_to_next_phase = transition_to_next_phase
        if simulated_cell_volume is None:
            self.simulated_cell_volume = 1
        else:
            self.simulated_cell_volume = simulated_cell_volume

        # the default rates are reference values for MCF-7, in 1/min
        if cytoplasm_volume_change_rate is None:
            self.cytoplasm_volume_change_rate = 0.27 / 60.0
        else:
            self.cytoplasm_volume_change_rate = cytoplasm_volume_change_rate
```

```
    if nuclear_volume_change_rate is None:
        self.nuclear_volume_change_rate = 0.33 / 60.0
    else:
        self.nuclear_volume_change_rate = nuclear_volume_change_rate
    if calcification_rate is None:
        self.calcification_rate = 0
    else:
        if calcification_rate < 0:
            raise ValueError(f"`calcification_rate` must be >= 0, got {calcification_rate}
                                                 ")
        self.calcification_rate = calcification_rate
    if fluid_change_rate is None:
        self.fluid_change_rate = 3.0 / 60.0
    else:
        self.fluid_change_rate = fluid_change_rate
    self.volume = CellVolumes(target_fluid_fraction=target_fluid_fraction,
        nuclear_fluid=nuclear_fluid, nuclear_solid=nuclear_solid,
        nuclear_solid_target=nuclear_solid_target,
        cytoplasm_fluid=cytoplasm_fluid, cytoplasm_solid=cytoplasm_solid,
        cytoplasm_solid_target=cytoplasm_solid_target,
        target_cytoplasm_to_nuclear_ratio=target_cytoplasm_to_nuclear_ratio,
        calcified_fraction=calcified_fraction,
        relative_rupture_volume=relative_rupture_volume)
```

### C Phase init function

```python
def __init__(self, target_fluid_fraction=None, nuclear_fluid=None, nuclear_solid=None,
    nuclear_solid_target=None,cytoplasm_fluid=None, cytoplasm_solid=None,
    cytoplasm_solid_target=None, target_cytoplasm_to_nuclear_ratio=None,
    calcified_fraction=None, relative_rupture_volume=None):
    _total = 2494
    _fluid_fraction = .75
    _fluid = _fluid_fraction * _total
    _solid = _total - _fluid
    _nuclear = 540
    _nuclear_fluid = _fluid_fraction * _nuclear
    _nuclear_solid = _nuclear - _nuclear_fluid
    _cytoplasm = _total - _nuclear
    _cytoplasm_fluid = _fluid_fraction * _cytoplasm
    _cytoplasm_solid = _cytoplasm - _cytoplasm_fluid
    _calcified_fraction = 0
    _relative_rupture_volume = 100
    if target_fluid_fraction is None:
        self.target_fluid_fraction = _fluid_fraction
    else:
        if not 0 <= target_fluid_fraction <= 1:
            raise ValueError(f"`target_fluid_fraction` must be in range [0, 1]. Got {
                                                                    target_fluid_fraction}")
        self.target_fluid_fraction = target_fluid_fraction
    if nuclear_fluid is None:
        self.nuclear_fluid = _nuclear * self.target_fluid_fraction
    else:
        if nuclear_fluid < 0:
            raise ValueError(f"`nuclear_fluid` must be >=0. Got {nuclear_fluid}")
        self.nuclear_fluid = nuclear_fluid
    if nuclear_solid is None:
        self.nuclear_solid = _nuclear * (1 - self.target_fluid_fraction)
    else:
        if nuclear_solid < 0:
            raise ValueError(f"`nuclear_solid` must be >=0. Got {nuclear_solid}")
        self.nuclear_solid = nuclear_solid
    if nuclear_solid_target is None:
        self.nuclear_solid_target = self.nuclear_solid
    else:
```

```
            if nuclear_solid_target < 0:
                raise ValueError(f"`nuclear_solid_target` must be >=0. Got {
                                                                                nuclear_solid_target}")
            self.nuclear_solid_target = nuclear_solid_target
        if cytoplasm_fluid is None:
            self.cytoplasm_fluid = _cytoplasm * self.target_fluid_fraction
        else:
            self.cytoplasm_fluid = cytoplasm_fluid
        if cytoplasm_solid is None:
            self.cytoplasm_solid = _cytoplasm * (1 - self.target_fluid_fraction)
        else:
            self.cytoplasm_solid = cytoplasm_solid
        if cytoplasm_solid_target is None:
            self.cytoplasm_solid_target = self.cytoplasm_solid
        else:
            self.cytoplasm_solid_target = cytoplasm_solid_target
        self.cytoplasm = self.cytoplasm_fluid + self.cytoplasm_solid
        self.nuclear = self.nuclear_fluid + self.nuclear_solid
        if target_cytoplasm_to_nuclear_ratio is None:
            self.target_cytoplasm_to_nuclear_ratio = self.cytoplasm / (1e-16 + self.
                                                                        nuclear)
        else:
            self.target_cytoplasm_to_nuclear_ratio = target_cytoplasm_to_nuclear_ratio
        if calcified_fraction is None:
            self.calcified_fraction = _calcified_fraction
        else:
            self.calcified_fraction = calcified_fraction
        if relative_rupture_volume is None:
            self.relative_rupture_volume = _relative_rupture_volume
        else:
            self.relative_rupture_volume = relative_rupture_volume
        self.fluid = self.cytoplasm_fluid + self.nuclear_fluid
        self.solid = self.cytoplasm_solid + self.nuclear_solid
        self.total = self.nuclear + self.cytoplasm
        self.fluid_fraction = self.fluid / self.total
        self.rupture_volume = self.relative_rupture_volume * self.total
```

#### D Ki-67 Basic Cycle Improved Division Implementation in CompuCell3D

Note: we have removed the implementation of population statistics, data saving, and plots from the code presented here. In places this would cause an error we have added a `pass` statement. The model that comes packaged with PhenoCellPy does include population statistics, data saving, and plots.

```
from cc3d.cpp.PlayerPython import *
from cc3d import CompuCellSetup
from cc3d.core.PySteppables import *
from numpy import median, quantile, nan
import sys
sys.path.extend(["C:\\PhenoCellPy"])
import Phenotypes as pheno
def Ki67pos_transition(*args):
    # print(len(args), print(args))
    # args = [cc3d cell volume, phase's target volume, time in phase, phase duration
    return args[0] >= args[1] and args[2] > args[3]

class ConstraintInitializerSteppable(SteppableBasePy):
    def __init__(self, frequency=1):
        SteppableBasePy.__init__(self, frequency)
        self.track_cell_level_scalar_attribute(field_name='phase_index_plus_1',
                                                             attribute_name='phase_index_plus_1')
        self.target_volume = None
        self.doubling_volume = None
        self.volume_conversion_unit = None

    def start(self):
        side = 10
        self.target_volume = side * side
        self.doubling_volume = 2 * self.target_volume
        x = self.dim.x // 2 - side // 2
        y = self.dim.x // 2 - side // 2
        cell = self.new_cell(self.CELL)
        self.cell_field[x:x + side, y:y + side, 0] = cell
        dt = 5  # 5 min/mcs
        ki67_basic_modified_transition = pheno.phenotypes.Ki67Basic(dt=dt,
                                                  transitions_to_next_phase=[None,
                                                                          Ki67pos_transition],
```

```python
                                             transitions_to_next_phase_args=[None,
                                                                              [-9, 1, -9, 1]])
        self.volume_conversion_unit = self.target_volume / \
            ki67_basic_modified_transition.current_phase.volume.total
        for cell in self.cell_list:
            cell.targetVolume = self.target_volume
            cell.lambdaVolume = 2.0
            pheno.utils.add_phenotype_to_CC3D_cell(cell, ki67_basic_modified_transition)
            cell.dict["phase_index_plus_1"] = \
                cell.dict["phenotype"].current_phase.index + 1
        self.shared_steppable_vars["constraints"] = self

class MitosisSteppable(MitosisSteppableBase):
    def __init__(self, frequency=1):
        MitosisSteppableBase.__init__(self, frequency)
        self.constraint_vars = None
        self.previous_number_cells = 0
        self.plot = True
        self.save = False
        if self.save:
            pass
    def start(self):
        self.constraint_vars = self.shared_steppable_vars["constraints"]
        if self.plot:
            pass
    def step(self, mcs):
        if not mcs and self.plot:
            pass
        elif not mcs % 50 and len(self.cell_list) - self.previous_number_cells > 0 \
          and self.plot:
            pass
        cells_to_divide = []
        n_zero = 0
        n_one = 0
        volumes = []
        time_spent_in_0 = []
        time_spent_in_1 = []
        for cell in self.cell_list:
            volumes.append(cell.volume)
            cell.dict["phenotype"].current_phase.simulated_cell_volume = cell.volume
            if cell.dict["phenotype"].current_phase.index == 0:
```

```
                n_zero += 1
                time_spent_in_0.append(cell.dict["phenotype"].current_phase.time_in_phase)
            elif cell.dict["phenotype"].current_phase.index == 1:
                n_one += 1
                time_spent_in_1.append(cell.dict["phenotype"].current_phase.time_in_phase)
                # args = [cc3d cell volume,
                #         doubling colume,
                #         time in phase,
                #         phase duration]
                args = [
                    cell.volume,
                    .9 * self.constraint_vars.doubling_volume,
                    # we use 90\% of the doubling volume because cc3d cells
                    # will always be slightly below their target due to the contact energy
                    cell.dict["phenotype"].current_phase.time_in_phase + \
                        cell.dict["phenotype"].dt,
                    cell.dict["phenotype"].current_phase.phase_duration]
                cell.dict["phenotype"].current_phase.transition_to_next_phase_args = args
            changed_phase, should_be_removed, divides = \
                cell.dict["phenotype"].time_step_phenotype()
            converted_volume = self.constraint_vars.volume_conversion_unit * \
                               cell.dict["phenotype"].current_phase.volume.total
            cell.targetVolume = converted_volume
            if changed_phase:
                cell.dict["phase_index_plus_1"] = \
                    cell.dict["phenotype"].current_phase.index + 1
            if divides:
                cells_to_divide.append(cell)
        if self.save or self.plot:
            pass
        for cell in cells_to_divide:
            # self.divide_cell_random_orientation(cell)
            # Other valid options
            # self.divide_cell_orientation_vector_based(cell,1,1,0)
            # self.divide_cell_along_major_axis(cell)
            self.divide_cell_along_minor_axis(cell)

    def update_attributes(self):
        # resetting target volume
        converted_volume = self.constraint_vars.volume_conversion_unit * \
                       self.parent_cell.dict["phenotype"].current_phase.volume.total
```

```
        self.parent_cell.targetVolume = converted_volume
        self.clone_parent_2_child()
        self.parent_cell.dict["phase_index_plus_1"] = \
            self.parent_cell.dict["phenotype"].current_phase.index + 1
        self.child_cell.dict["phase_index_plus_1"] = \
            self.child_cell.dict["phenotype"].current_phase.index + 1
        self.child_cell.dict["phenotype"].time_in_phenotype = 0
    def on_stop(self):
        self.finish()
    def finish(self):
        if self.save:
            pass
```

### E Ki-67 Basic Cycle Implementation in Tissue Forge

```
import tissue_forge as tf
import numpy as np
import sys
sys.path.extend(["C:\\PhenoCellPy"])
import Phenotypes as pheno
def get_radius_sphere(volume):
    return ((1 / (np.pi * 4 / 3)) * volume) ** (1 / 3)
# potential cutoff distance
cutoff = 3
# space set up
dim = [50, 50, 50]
tf.init(dim=dim, cutoff=cutoff)
pot = tf.Potential.morse(d=3, a=5, min=-0.8, max=2)
# Particle types
mass = 40
radius = .4
global density
density = mass / ((4 / 3) * np.pi * radius * radius * radius)
dt = 10  # min/time step
ki67_basic = pheno.phenotypes.Ki67Basic(dt=dt)
global volume_conversion_unit
volume_conversion_unit = mass/ki67_basic.current_phase.volume.total
class CellType(tf.ParticleTypeSpec):
    mass = mass
    target_temperature = 0
    radius = radius
    dynamics = tf.Overdamped
    cycle = ki67_basic
Cell = CellType.get()
tf.bind.types(pot, Cell, Cell)
rforce = tf.Force.random(mean=0, std=50)
# bind it just like any other force
tf.bind.force(rforce, Cell)
first_cell = Cell([d // 2 for d in dim])
first_cell.cycle = ki67_basic
global cells_cycles
cells_cycles = {f"{first_cell.id}": ki67_basic}
def step_cycle_and_divide(event):
```

```
    for p in Cell.items():
        pcycle = cells_cycles[f"{p.id}"]
        pcycle.current_phase.simulated_cell_volume = p.mass * density
        phase_change, should_be_removed, division = pcycle.time_step_phenotype()
        radius = get_radius_sphere(
                volume_conversion_unit*pcycle.current_phase.volume.total)
        # book-keeping, making sure the simulated cell grows
        p.radius = radius
        p.mass = ((4 / 3) * np.pi * radius * radius * radius) * density
        # if division occurs, divide
        if division:
            print("@@@\nDIVISION\n@@@")
            # save cell attribs to halve later
            cur_mass = p.mass
            # divide and reasign attribs (is this step necessary?)
            child = p.split()
            cells_cycles[f"{child.id}"] = ki67_basic
            child.mass = p.mass = cur_mass / 2
            child.radius = p.radius = get_radius_sphere((cur_mass / 2) / density)
            cells_cycles[f"{child.id}"].volume = child.mass * density
            cells_cycles[f"{child.id}"].simulated_cell_volume = child.mass * density
    return 0

tf.event.on_time(invoke_method=step_cycle_and_divide, period=.9*tf.Universe.dt)
# run the simulator interactive
tf.run()
```
